# Similar metabolic pathways are affected in both Congenital Myasthenic Syndrome-22 and Prader-Willi Syndrome

**DOI:** 10.1101/2023.11.12.566752

**Authors:** Kritika Bhalla, Karen Rosier, Yenthe Monnens, Sandra Meulemans, Ellen Vervoort, Lieven Thorrez, Patrizia Agostinis, Daniel T. Meier, Anne Rochtus, James L Resnick, John W.M. Creemers

## Abstract

Loss of prolyl endopeptidase-like (*PREPL*) encoding a serine hydrolase with (thio)esterase activity leads to the recessive metabolic disorder Congenital Myasthenic Syndrome-22 (CMS22). It is characterized by severe neonatal hypotonia, feeding problems, growth retardation, and hyperphagia leading to rapid weight gain later in childhood. The phenotypic similarities with Prader-Willi syndrome (PWS) are striking, suggesting that similar pathways are affected. The aim of this study was to identify changes in the hypothalamic-pituitary axis in mouse models for both disorders and to examine mitochondrial function in skin fibroblasts of patients and knockout cell lines. We have demonstrated that *Prepl* is downregulated in the brains of neonatal PWS-IC^-p/+m^ mice. In addition, the hypothalamic-pituitary axis is similarly affected in both *Prepl*^-/-^ and PWS-IC^-p/+m^ mice resulting in defective orexigenic signaling and growth retardation. Furthermore, we demonstrated that mitochondrial function is altered in *PREPL* knockout HEK293T cells and can be rescued with the supplementation of coenzyme Q10. Finally, PREPL-deficient and PWS patient skin fibroblasts display defective mitochondrial bioenergetics. The mitochondrial dysfunction in PWS fibroblasts can be rescued by overexpression of PREPL. In conclusion, we provide the first molecular links between CMS22 and PWS, raising the possibility that PREPL substrates might become therapeutic targets for treating both disorders.

**Graphical Abstract:** 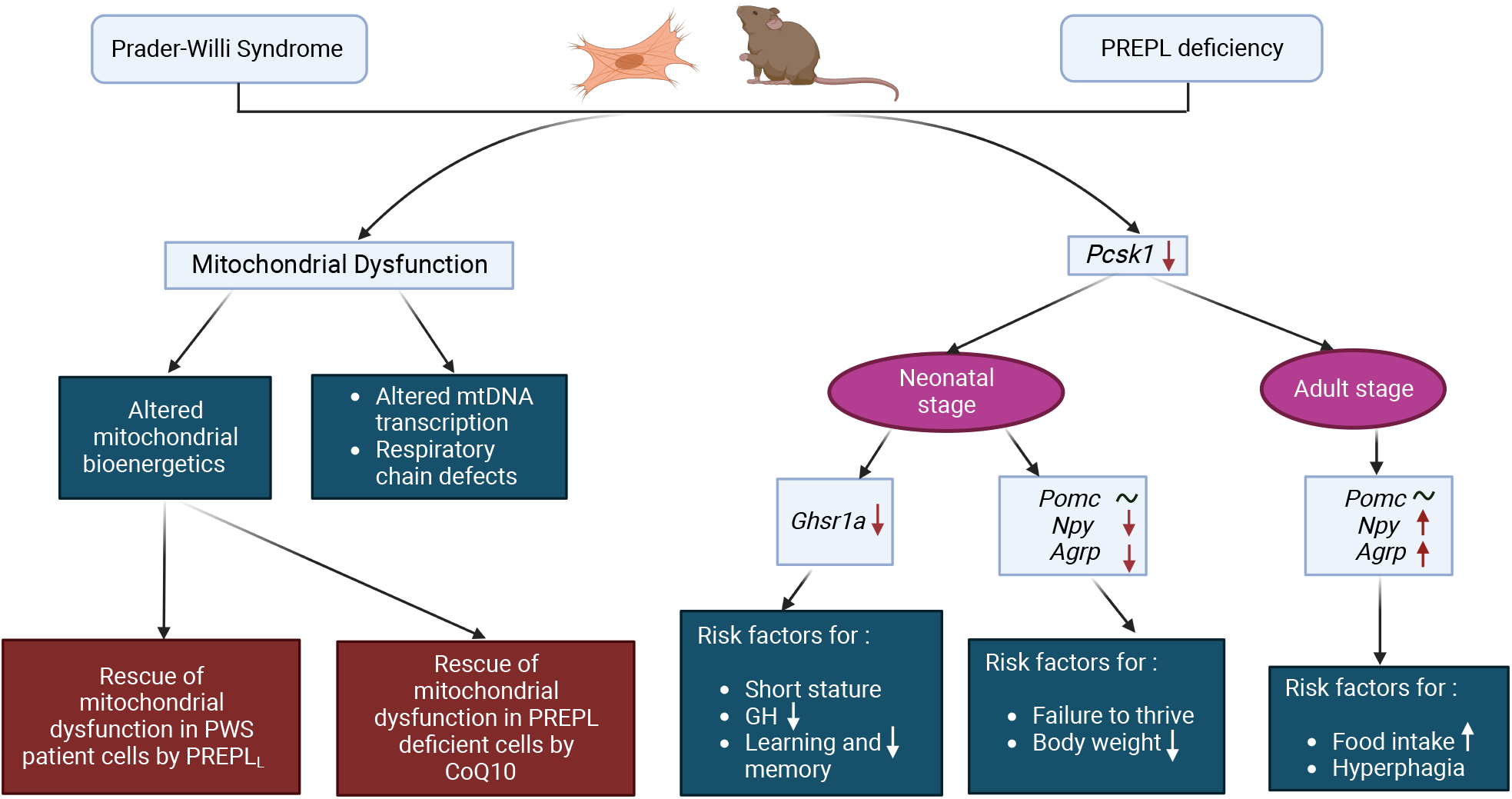

## Introduction

Mutation in the prolyl endopeptidase-like (*PREPL*) gene on the 2p21 chromosome leads to the recessive metabolic disorder called Congenital Myasthenic Syndrome-22 (CMS22; OMIM 616224) (1, 2). This syndrome is characterized by neonatal hypotonia, growth hormone (GH) deficiency, cognitive impairment and feeding problems leading to failure to thrive in early childhood, followed by hyperphagia and rapid weight gain during adolescence. The hypotonia spontaneously improves, but facial weakness, eyelid ptosis, nasal voice, and mild proximal muscle weakness persist. In addition, a dry mouth with thickened saliva and hypergonadotropic hypogonadism are frequently observed. However, the molecular mechanisms by which PREPL deficiency leads to these clinical phenotypes are currently unknown.

PREPL deficiency shares striking clinical and pathophysiological similarities with another neurodevelopmental disorder called Prader-Willi Syndrome (PWS). This rare disorder is caused by the loss of paternal expression of genes on the maternally imprinted chromosome 15q11-13. PWS patients also have neonatal hypotonia, failure to thrive, feeding difficulties that advance towards hyperphagia and severe childhood obesity, GH-deficiency, and cognitive impairment. The dramatic switch in phenotype from failure to thrive to hyperphagia on the one hand, and the spontaneous improvement of the hypotonia during the first years of life on the other, is uniquely linked to these two syndromes (3). Consequently, most PREPL-deficient patients are initially diagnosed with PWS phenotype without PWS genotype (1). On the other hand, PWS patients have several additional phenotypes that are not observed in CMS22, such as small hands and feet, behavioral problems, and skin picking. This is not surprising since PWS results from the loss of expression of multiple genes located at 15q11-q13 genetic locus.

In 2017, Burnett et al. observed reduced expression levels of proprotein convertase 1/3 (PC1/3), encoded by *PCSK1*, and its potential upstream regulator, nescient helix-loop-helix 2 (*NHLH2*) in induced pluripotent stem cell-derived neurons of PWS patients and hypothalami of Snord116^-p/+m^ mouse model (4). However, Polex-Wolf et al. have contested this finding (5). A recent study by Duarte et al. revealed that the expression of PC1/3 is reduced in *Prepl* knockdown mouse neuroblastoma cells (2). PC1/3 is involved in the cleavage and hence activation of precursor proteins into active hormones, growth factors and neuropeptides (6). Some of these factors are known to regulate food intake, body weight and growth, e.g., pro-opiomelanocortin (*POMC*), pro-neuropeptide Y (*proNPY*), pro-agouti-related peptide (*proAGRP*), pro-growth hormone-releasing hormone (*proGHRH*), vasopressin and oxytocin (7, 8). PC1/3 deficient mice and humans display malabsorptive diarrhea, hypogonadism, decreased GH and hyperphagia resulting in obesity (4, 9). PC1/3 expression and proproteins affecting leptin/melanocortin signalling have not been studied in the context of PREPL deficiency.

PREPL is a ubiquitously expressed serine hydrolase having *in vitro* (thio)esterase activity (10). Its highest expression is found in the brain, while intermediate levels are detected in skeletal muscle, heart, and kidney. The most abundant form of PREPL is a soluble cytosolic protein partly colocalizing with cytoskeletal elements, Golgi apparatus, growth cones, and vesicular structures (11). Recently, PREPL has also been identified as a mitochondrial protein (10). These two isoforms are encoded by different splice variants: PREPL_L_ (long; 727 amino acids) and PREPL_S_ (short; 638 amino acids), the former residing in mitochondria by virtue of an amino-terminal mitochondrial targeting signal that is not present in PREPL_S_. PREPL_S_, the cytoplasmic form, is involved in regulated secretion as revealed by electrophysiological studies at the neuromuscular junction (NMJ), AP-1 binding, and *in vitro* secretion studies (2, 12). The interactome of PREPL contains a considerable number of mitochondrial proteins which are mostly involved in oxidative phosphorylation (OXPHOS) and mitochondrial translation (10).

Recently, we and others described the process whereby translation of mitochondrial-DNA (mt-DNA) encoded respiratory chain complex subunits is impaired in *Prepl*^-/-^ mice (10, 13). Moreover, isolated complex IV and combined respiratory chain complex deficiencies have also been described in several PREPL-deficient patients and mice (1, 14–17). Apart from this, electron micrographs of mitochondria revealed structural abnormalities at the nerve terminal of a PREPL-deficient patient and quadriceps of *Prepl*^-/-^ mice. Mitochondria generate energy and maintain Ca^2+^ homeostasis. Mitochondrial dysfunction can result in several phenotypes like skeletal muscle weakness, ptosis, failure to thrive, fatigue, reduced metabolic rate, decreased energy, learning difficulties, developmental delay, obesity, seizures, and stroke, many of which are found in CMS22 and PWS (18–20). The similarities in phenotypes between CMS22 and PWS have raised the question of whether the same or overlapping pathways are affected (21, 22). Recently, Butler et al. suggested that disturbances in cellular metabolism and mitochondrial bioenergetics contribute to the pathophysiology of PWS (23). Consistently, oxidative phosphorylation complex activities have recently been shown to be impaired in PWS mice (24). Taken together, exploring the mitochondrial function might reveal similarly affected pathways and help understand the pathophysiology of PREPL deficiency and PWS.

Supplementation with coenzyme Q_10_ (CoQ_10_), an electron carrier of the mitochondrial electron transport chain (ETC), often has therapeutic benefits for patients with respiratory chain defects (25). CoQ_10_ deficiency presents phenotypes including central nervous system dysfunction and muscle weakness, partially overlapping with PWS and CMS22 patients (26). Moreover, plasma CoQ_10_ levels are often reduced in PWS patients compared to aged-matched non-obese patients (27). A study of improved psychomotor development has been reported in PWS infants following supplementation with CoQ_10_ and GH therapy (28). A clinical trial led by Dr. Ingrid Tein (Hospital for Sick Children, Toronto, Canada) is currently in progress to determine the possible benefits of CoQ_10_ supplementation in PWS patients. Patients with mitochondrial disorders are frequently treated with CoQ_10_ supplements to improve muscle dysfunction. However, efficacy in improving neurological symptoms remains variable and inconclusive.

In this paper, we have studied similarities in the pathophysiology of CMS22 and PWS. Using mouse models for PREPL deficiency and PWS we have examined the leptin/melanocortin signalling pathway. Furthermore, we have performed mitochondrial respiration assays using genetically modified cell lines and fibroblasts from both PREPL-deficient and PWS patients. Moreover, we have studied the effect of coenzyme Q_10_ (CoQ_10_) supplementation on mitochondrial function in the absence of PREPL. Finally, we have investigated the potential rescue of cellular respiration and mitochondrial bioenergetics in PWS patient cell lines by overexpressing PREPL_L_.

## Results

### 1.1. *Prepl* expression is reduced in the brain of PWS-IC^-p/+m^ mice

Figure 1A is a schematic representation of the PWS locus. In PWS-IC^-p/+m^ mice, a 35 kb region including the imprinting center (IC) and exons 1-6 of the *Snurf/Snrpn* gene are deleted (29, 30). This was confirmed by the lack of expression of *Snrpn* using RT-qPCR (Figure 1B). To explore a potential molecular link between PREPL deficiency and PWS, we examined *Prepl* expression levels in whole brains of neonatal PWS^-p/+m^ mice. A prominent 2-fold reduction of *Prepl* expression was observed in the brains of neonatal PWS-IC^-p/+m^ mice (Figure 1C-E). The primers used in Figure 1C and D detect both the long isoform (mitochondrial) and short isoform (cytoplasmic) of *Prepl*, while the primers in Figure 1E are specific to the long form. Both the transcript levels of the long and the short isoforms were significantly reduced. As expected, no expression of *Prepl* was detected in *Prepl*^-/-^ mice in Figure 1C since the primers were directed against the region encoded by the deleted exon 10. The reduced expression of *Prepl_L_* observed in *Prepl*^-/-^ mice (Figure 1D) is most likely the consequence of nonsense-mediated mRNA decay due to the frameshift caused by exon 10 deletion. The data obtained in Figure 1C-E were obtained from two different cohorts of PWS-IC^-p/+m^ mice (shown separately in Supplementary figure 1). Remarkably, we noticed a significant reduction of *Prepl* expression in the first cohort compared to the second, despite using identical procedures. Lack of expression of *Snord116*, a small nucleolar RNA (snoRNA) leads to many neuroendocrine phenotypes observed in both PWS humans and mice, making Snord116^-p/+m^ mutant mice a valuable model for studying PWS (31). Hence, we investigated transcript levels of *Prepl* in the whole brains of neonatal Snord116^-p/+m^ mice. However, they remained unaltered compared to the controls (Figure 1F-H).

**Figure 1:**
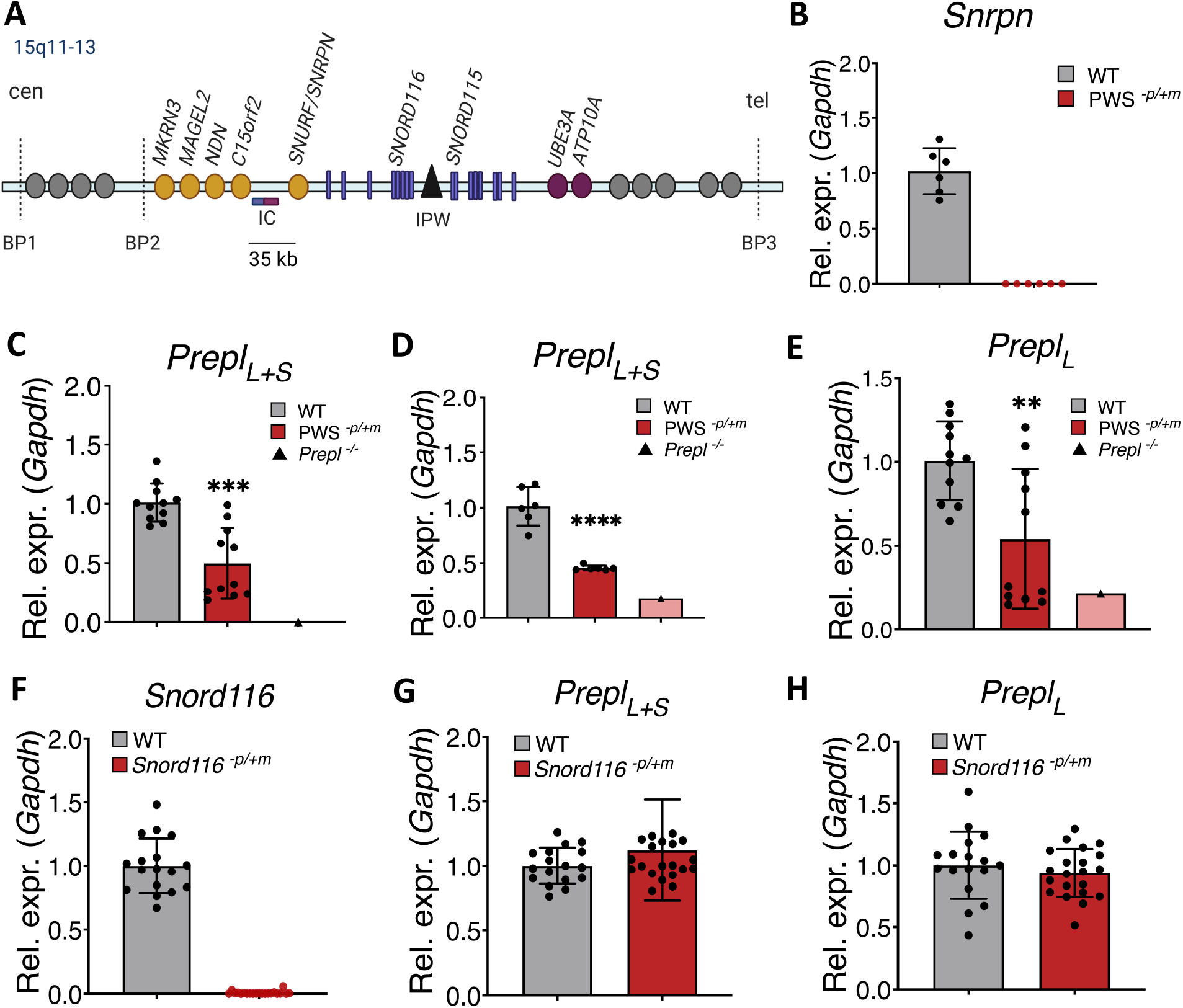
*Prepl* is downregulated in the brain of newborn PWS-IC*^-p/+m^* mice. (**A**) Diagram of the PWS genetic locus. Paternally expressed genes are shown in yellow, maternally expressed genes in magenta and non-imprinted genes in gray. Protein-coding genes are shown in ovals, snoRNAs as purple rectangles, long noncoding RNAs as black triangles and imprinting center as purple-magenta rectangle. Not drawn to scale. Cen, centromere; tel, telomere. The 35 kb deleted region in PWS-IC*^-p/+m^* mice is underlined. (**B**) *Snrpn* in the PWS locus is not expressed in PWS-IC*^-p/+m^* mice, as measured by RT-qPCR. (**C** and **D**) Reduced gene expression levels of *Prepl_L+S_* in the whole brains of PWS-IC*^-p/+m^* mice. The primer pairs are directed towards exon 10 and exon 7/8 of *Prepl* respectively. (**E**) Altered transcript levels of *Prepl_L_* in the brains of PWS-IC*^-p/+m^* mice. The primer pair is directed towards exon 2/3 of *Prepl*, present in the long but not the short isoform. (**F**) Expression of *Snord116* in whole brains of Snord116*^-p/+m^* mice. (**G** and **H**) Transcript levels of *Prepl* (long and short isoforms) remain unchanged in Snord116*^-p/+m^* mouse model. Each data point represents 1 animal (N > 6 per genotype; N = 1 *Prepl*^-/-^), plotted as mean ± SD. Differences to the unaffected controls were either analyzed with a 2-tailed, type 3 (assumes unequal variance) Student’s t-test (**B, F, G, H**) or one-way ANOVA with Tukey’s test for multiple comparisons (**C, D, E**). *p < 0.05, **p < 0.01, ***p < 0.001, ****p < 0.0001.

### 1.2. *Pcsk1*, *Npy*, and *Ghsr1a* are downregulated in the whole brains of newborn *Prepl*^-/-^ and PWS-IC^-p/+m^ mice

We subsequently studied the expression of *Pcsk1* and its proposed upstream regulator, the transcription factor *Nhlh2* in the whole brain of newborn *Prepl*^-/-^ and PWS-IC^-p/+m^ mice (4). In both models, *Pcsk1* was significantly downregulated, albeit much more pronounced in the PWS-IC^-p/+m^ mice (Figure 2A (p= 0.0003, PWS-IC^-p/+m^); Figure 2B (p= 0.03, *Prepl*^-/-^)). *Nhlh2* expression levels were upregulated in both *Prepl^-/-^* (not significant) and PWS-IC^-p/+m^ pups in whole brain tissue (Figure 2C, D), in contrast to the published downregulation in PWS (4). Since both *Prepl*^-/-^ and PWS-IC^-p/+m^ mutant mice have postnatal feeding problems and growth retardation (10, 30), we assessed the expression levels of orexigenic and anorexigenic neuropeptides as well as the growth hormone secretagogue receptor (*Ghsr1a*). The expression levels of the primary appetite-stimulating neuropeptide Y (*Npy*) were significantly reduced in newborn pups of both models (Figure 2E, F). mRNA levels of agouti-related peptide (*Agrp*) were decreased in PWS-IC^-p/+m^ mice but remained unaltered in *Prepl*^-/-^ mice (Figure 2G, H). Levels of Pro-opiomelanocortin (*Pomc*) neuropeptide, the precursor to the satiety factor melanocyte-stimulating hormone, remained unchanged in the brains of both *Prepl*^-/-^ and PWS-IC^-p/+m^ mice (Figure 2I, J). The expression levels of *Ghsr1a*, also called ghrelin receptor were strongly downregulated in the brain tissues of PWS-IC^-p/+m^ and *Prepl*^-/-^ mice (Figure 2K, L). The gene expression levels of *Pcsk1,* orexigenic neuropeptide *Agrp* and *Ghsr1a* were not affected in the whole brains of newborn Snord116^-p/+m^ (Supplementary figure 2). Taken together, the leptin/melanocortin pathway is similarly affected in both PWS-IC^-p/+m^ and *Prepl*^-/-^ models albeit more severely in PWS-IC^-p/+m^ mice.

**Figure 2:**
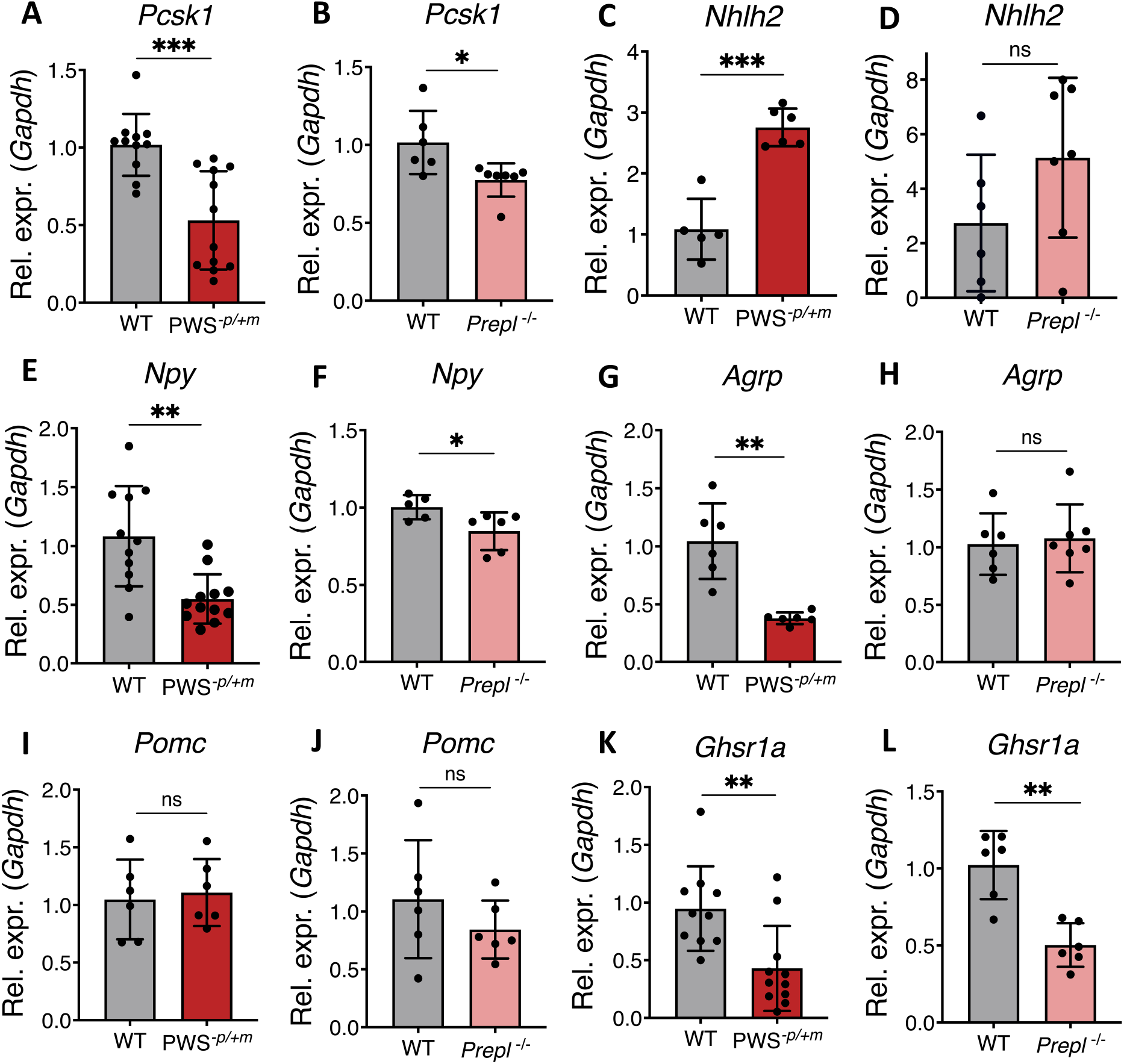
The leptin/melanocortin pathway is altered in both PWS-IC*^-p/+m^* and *Prepl*^-/-^ mouse models. (**A** and **B**) Reduced gene expression levels of proprotein convertase subtilisin/kexin type 1 (*Pcsk1*) in whole brains of neonatal PWS-IC*^-p/+m^* and *Prepl*^-/-^ mice. (**C** and **D**) Increased transcript levels of nescient helix loop helix 2 (*Nhlh2)* in both mouse models. (**E-H**) mRNA expression of neuropeptide Y (*Npy*) and Agouti-related peptide (*Agrp*) in the newborn PWS-IC*^-p/+m^* and *Prepl*^-/-^ mice. (**I** and **J**) Unaltered levels of pro-opiomelanocortin (*Pomc*) in both PWS-IC*^-p/+m^* and *Prepl*^-/-^ mice. (**K** and **L**) Downregulation of growth hormone secretagogue receptor 1a (*Ghsr1a*) in PWS-IC*^-p/+m^* and *Prepl*^-/-^ mice. Each data point represents 1 animal (N > 5 per genotype), plotted as mean ± SD. Differences to the unaffected controls were analyzed using a 2-tailed, type 3 (assumes unequal variance) Student’s t-test. *p < 0.05, **p < 0.01, ***p < 0.001.

Hyperphagia, starting during childhood, is one of the symptoms observed in both CMS22 and PWS. Therefore, we measured the relative and absolute food intake of young adult (8-weeks-old) *Prepl*^-/-^ mice. The absolute food intake was unaltered, but when divided by body weight to compensate for the smaller size of *Prepl*^-/-^ mice, the food intake of *Prepl*^-/-^ was mildly but significantly increased compared to wild-type controls (Figure 3A, B). Subsequently, the *Pcsk1* expression levels were studied in adult *Prepl*^-/-^ mice after overnight fasting or overnight fasting and 5h refeeding. A non-significant reduction of *Pcsk1* was observed in the hypothalami of fasting mice (p=0.0542) but remained unchanged after refeeding (Figure 3C). The *Nhlh2* and *Pomc* expression was not significantly decreased in fasting or refed conditions in the *Prepl*^-/-^ mice Figure 3D, E). Surprisingly, expression levels of the orexigenic factors *Npy* and *Agrp* were significantly increased in *Prepl*^-/-^ mice after refeeding, suggesting disturbed satiety (Figure 3F and G). Lastly, the expression levels of *Snord116* remained unaltered in *Prepl^-^*^/-^ mice, independent of the nutritional state (Figure 3H). The last model system we used to study *Pcsk1* expression was *Prepl* KO βTC3 insulinoma cells. These neuroendocrine cells with high endogenous PREPL levels are suitable for studying the regulated secretion of large dense-core vesicles. These PREPL-deficient cells revealed a significant reduction of *Pcsk1* expression (KO1: 35% decrease, KO2: 45% decrease, KO3: 34% decrease) (Figure 3I) and ∼40% reduction in PC1/3 protein expression in these cell lines (Figure 3J).

**Figure 3:**
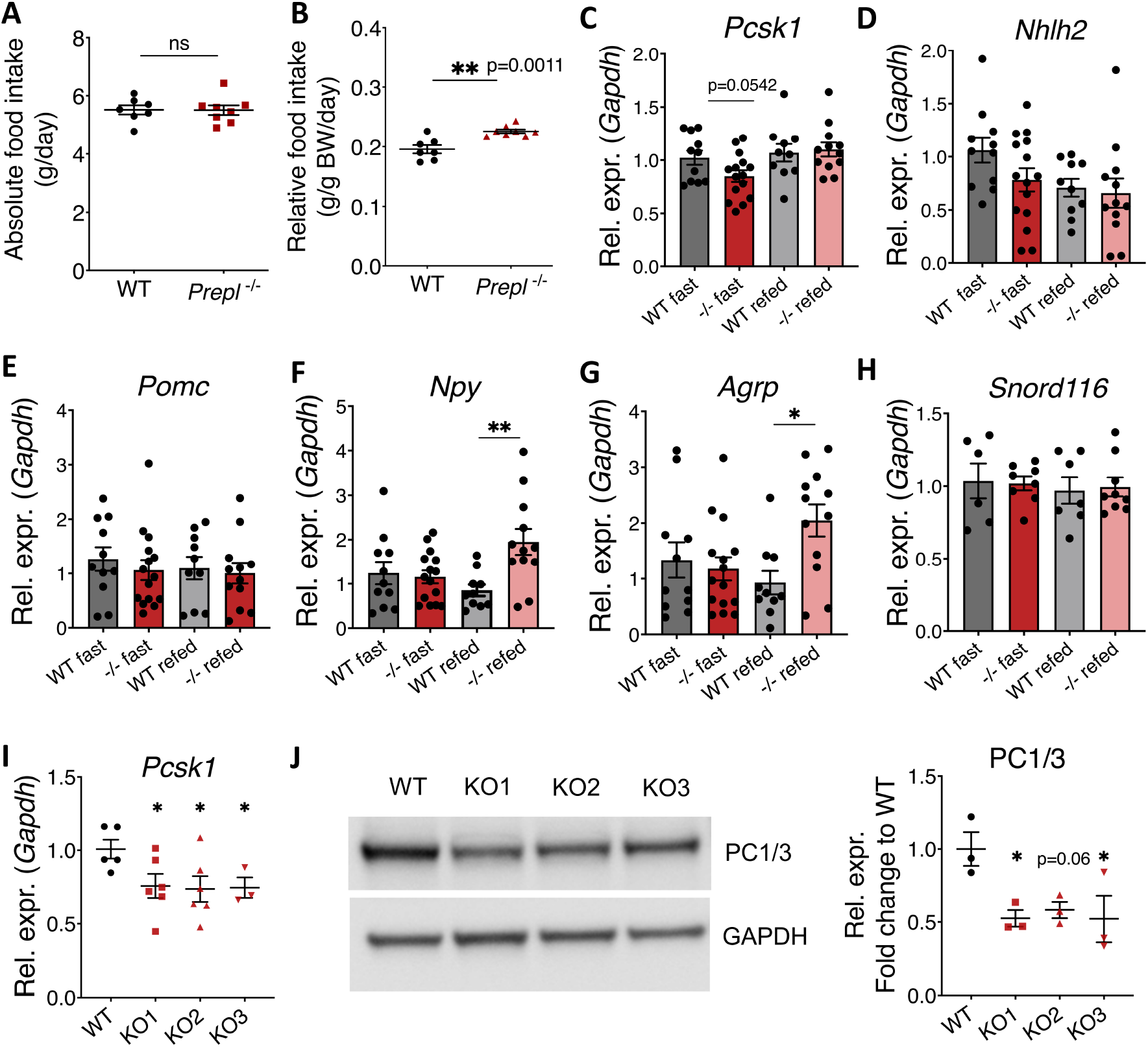
Hypothalamic loss of *Prepl* results in dysfunctional orexigenic signaling and mild hyperphagia in adult mice. (**A**) Food intake of 8-week-old in *Prepl^-/-^* male mice and wildtype littermates (N=7-8). (**B**) Relative food intake, normalized to body weight. Data are represented as mean ± SD. Differences to the controls were analyzed using unpaired Student’s t-test. **p < 0.01. (**C** and **D**) Transcript levels of hypothalamic *Pcsk1* and *Nhlh2* after 16h fasting or 16h fasting + 5h refeeding). (**E**) Expression levels of anorexigenic neuropeptide *Pomc* in *Prepl^-/-^*mice. (**F** and **G**) Expression levels of the orexigenic neuropeptides, *Npy* and *Agrp* (**H**) Expression levels of *Snord116*. Transcript levels were analyzed by RT-qPCR in 12-week-old male mice (N=10-15). Expression levels were normalized to *Gapdh.* Data are represented as mean ± SD and were analyzed using One-way ANOVA with Dunnett’s multiple comparison test. For *Pcsk1* an unpaired Student’s t-test was performed to compare the WT and KO fast condition. * p<0.05, ** p<0.01. (**I**) Quantification of *Pcsk1* expression levels in three different *Prepl* KO βTC3 cells lines. (**J**) Protein levels of PC1/3 are decreased in *Prepl* KO βTC3 cells. Representative quantification of three independent experiments. Data are represented as mean ± SD (N=3), analyzed using one-way ANOVA with Dunnett’s multiple comparisons test. * p<0.05.

### 1.3. PREPL deficiency results in impaired mitochondrial function in HEK293T cells

Neurons critically depend on mitochondrial function, not only for establishing membrane excitability but also for neurotransmission and plasticity (32). Since PREPL_L_ resides in mitochondria and several CMS22 patients have impaired mitochondrial function, we decided to measure the functional capacity of mitochondria in the absence of PREPL. The real-time measurements represented as oxygen consumption rates (OCRs) were initially performed on wild type and *PREPL* KO HEK293T cells using Seahorse XFp extracellular flux analyzer. The overall OCR was significantly reduced in *PREPL* KO HEK293T cells (Figure 4A). Based on OCR, the contribution of individual metabolic parameters like basal respiration (substrate utilization), ATP-linked respiration, maximal respiratory capacity, spare capacity and non-mitochondrial respiration were calculated. Basal respiration, defined as the net energetic demand of cells under baseline conditions was considerably decreased in *PREPL* KO HEK293T cells (Figure 4B). Next oligomycin, an ATP synthase (complex V) inhibitor, was added. The decrease in OCR corresponds to the portion of basal respiration used to drive ATP production (33). Figure 4C shows that ATP-linked respiration was significantly reduced in *PREPL* KO HEK293T cells. Subsequently, the cells were incubated with the uncoupler FCCP, making the inner mitochondrial membrane permeable to protons, thereby disrupting the proton gradient and membrane potential. As a result, the electron flow through the electron transport chain (ETC) becomes uninhibited, allowing the cells to consume oxygen at their maximal bio-energetic limit. This maximal capacity was also significantly decreased in the absence of PREPL (Figure 4D). A final series of measurements were performed after the addition of antimycin A, an inhibitor of complex III. It allows the measurement of non-mitochondrial respiration, which was also decreased suggesting additional dysfunctional oxidative pathways (Figure 4E). The spare respiratory capacity (Figure 4F) of cells, defined as the difference between maximal and basal respiration was also significantly reduced in *PREPL* KO cells. This indicates a decreased ability of cells to respond to additional energy demands under stress. The apparent proton leak is a phenomenon in which protons or H+ ions are transported into the mitochondrial matrix without contributing to the proton motive force required to produce ATP. The apparent proton leak was reduced in *PREPL* KO cells compared to WT cells (Figure 4G). Overall, this data confirms that mitochondrial performance is disturbed in the absence of PREPL.

**Figure 4:**
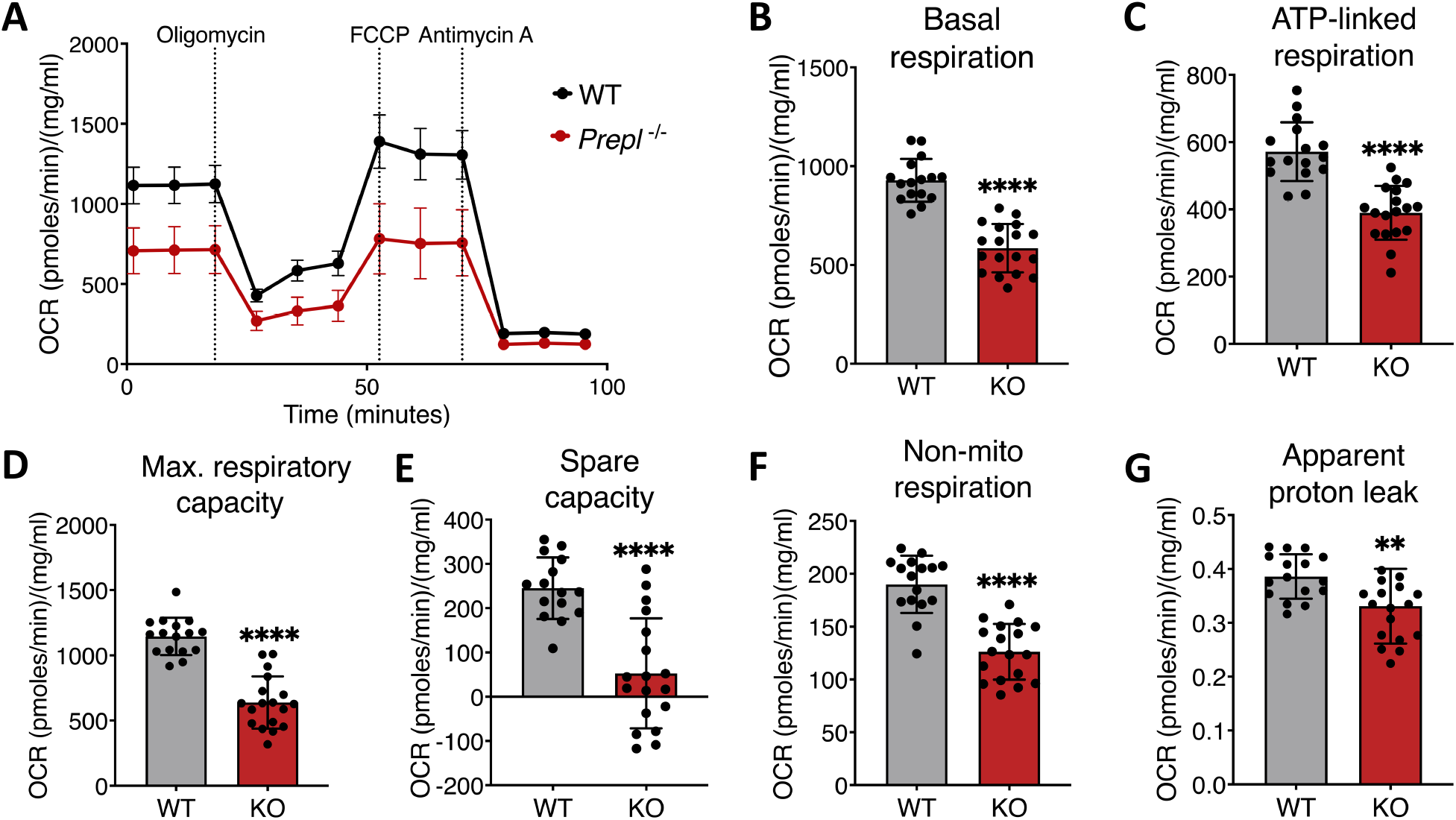
Loss of PREPL impairs mitochondrial function in HEK293T cells. (**A**) Seahorse respiratory flux profiles indicate that the oxygen consumption rate (OCR) is reduced in *PREPL* KO HEK293T cells compared to the control (WT). Bioenergetic parameters (**B**) Basal respiration, (**C**) ATP- linked or coupled respiration (**D**) Maximal (Max.) respiratory capacity were significantly reduced in *PREPL* KO compared to the control cells. (**E**) Non-mitochondrial (mito) respiration adjustment (**F**) Spare reserve capacity (**G**) Apparent proton leak. Each data point represents an OCR measurement, plotted as mean ± SD. N > 15 wells for three independent experiments. Differences to the controls were analyzed using a 2-tailed Student’s t-test. *p < 0.05, **p < 0.01, ***p < 0.001, ****p < 0.0001. All the OCR measurements were normalized to the protein concentration.

### 1.4. PREPL deficient and PWS patient cell lines have altered cellular respiration and mitochondrial dysfunction

Since mitochondrial dysfunction has also been reported in PWS patients, we subsequently determined the functional capacity of mitochondria in skin fibroblasts obtained from PWS and PREPL-deficient patients (Figure 5A-G). A similar reduction in OCR was observed in both PWS and PREPL-deficient patient cell lines (Figure 5A) indicating abnormalities in the energy balance. Basal respiration and ATP-linked respiration were significantly affected in both PWS and PREPL-deficient fibroblasts (Figure 5B and C). Maximal respiratory capacity was notably reduced in PWS patients and shows a trend of reduction (p= 0.0783) in the PREPL-deficient patients (Figure D). Spare capacity, non-mitochondrial respiration, and apparent proton leak remain unchanged (Figure 5E, F and G). Taken together, these data demonstrate that mitochondrial dysfunction is found in both PREPL-deficient and PWS patient fibroblast cells.

**Figure 5:**
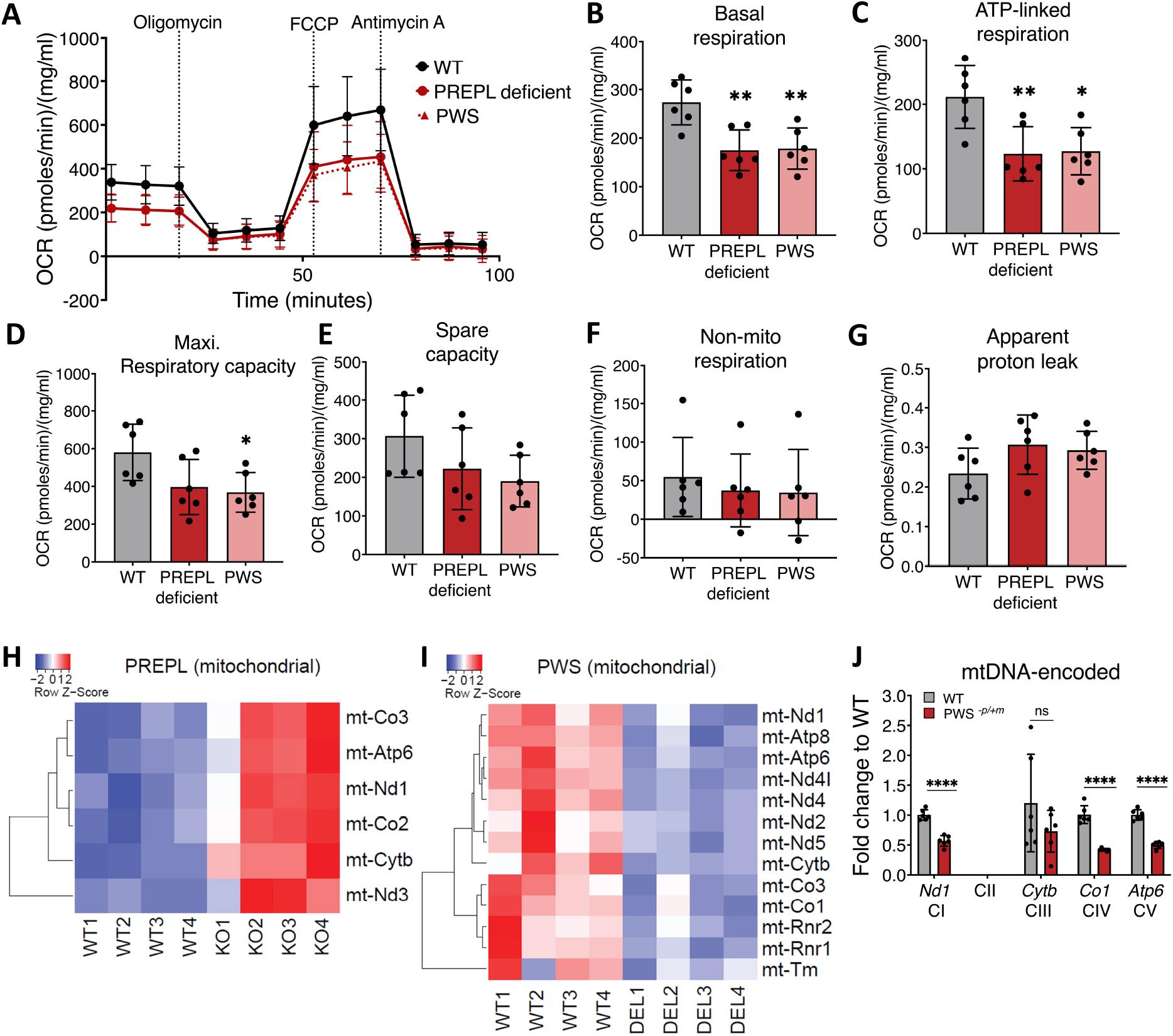
Mitochondrial dysfunction and altered cellular respiration in fibroblasts of PWS and PREPL deficient patients. (**A**) XFp analysis of cell mitochondrial stress shows reduced OCR levels for patients with PWS and PREPL deficiency compared to the controls. Bioenergetic parameters (**B** and **C**) Basal respiration, ATP-linked or coupled respiration adjustment were significantly reduced in PWS and PREPL deficient patients (**D**) Maximal (Max.) respiratory capacity is reduced in PWS and PREPL deficient patient fibroblasts. (**E**) Spare reserve capacity (**F**) Non-mitochondrial (mito) respiration and (**G**) Apparent proton leak remains unchanged in PWS and PREPL deficient patients. Each data point represents an OCR measurement, plotted as mean ± SD. N > 30 wells for 3 independent experiments for two WT, two PWS, and two PREPL deficient patients. Differences to the controls were analyzed using one-way ANOVA, with Tukey’s test for multiple comparisons. *p < 0.05, **p < 0.01. All the OCR measurements were normalized to the protein concentration. (**H**) Heatmap displaying differentially expressed mitochondrial genes in *Prepl*^-/-^ mice. (**I**) Heatmap displaying differentially expressed mitochondrial genes in PWS-IC*^-p/+m^* mice. (**J**) Decreased transcription of mitochondrial DNA (mtDNA) encoded complex subunits in PWS-IC*^-p/+m^* mice. Gene expression was analyzed by RT-qPCR in brains of newborn PWS-IC*^-p/+m^* mice (N=6 per genotype). *Gapdh* was used for normalization. **p < 0.01, ****p < 0.0001.

To investigate these pathways in our *in vivo* models, we performed RNA sequencing (RNAseq) on whole brains of neonatal *Prepl*^-/-^ and PWS-IC^-p/+m^ mice. The heat maps in Figure 5H and 5I depict differentially regulated mitochondrially linked genes (including those involved in mitochondrial respiration) in *Prepl*^-/-^ and PWS-IC^-p/+m^ mice compared to their WT littermates (p<0.1). As previously reported, gene expression of mtDNA-encoded subunits was upregulated in *Prepl*^-/-^ mice, while protein levels were reduced (10). A role for PREPL in mitochondrial protein synthesis was recently confirmed by Kramer *et al*. (13). The expression of mitochondrial-linked genes was downregulated in PWS-IC^-p/+m^ mice compared to their controls. To confirm the RNAseq data, we measured transcript levels of OXPHOS complex subunits in the brains of neonatal PWS-IC^-p/+m^ mice by RT-qPCR. As expected, steady-state levels of mitochondrial DNA-encoded (mtDNA) complex subunits were decreased in the brains of PWS-IC^-p/+m^ mice (Figure 5J). The expression levels of cytochrome b, the mtDNA-encoded subunit of complex III, were not significantly affected (Figure 5J). The steady-state levels of nuclear DNA-encoded mitochondrial subunits were also significantly reduced in PWS-IC^-p/+m^ mice (Supplementary figure 5A). Lastly, RT-qPCR revealed altered mitoribosome function in the whole brains of PWS-IC^-p/+m^ mice, indicating impaired protein translation and respiratory chain defects (Supplementary figure 5B, C). Taken together, these data show that protein synthesis in both *Prepl*^-/-^ and PWS-IC^-p/+m^ mice is downregulated, albeit through different mechanisms.

### 1.5. CoQ_10_ supplementation rescues the functional capacity of mitochondria in PREPL-deficient HEK293T cells

Since treatment of PWS patients with CoQ_10_ improves psychomotor development (28), we tested whether treatment of PREPL-deficient HEK293T cells for 5 days with 5µM CoQ_10_ also resulted in improvement of the mitochondrial function (Figure 6). OCRs were progressively increased, and metabolic parameters linked to the respiratory profile were significantly boosted in PREPL-deficient HEK293T cells following the treatment (Figure 6A). Defects in basal respiration, ATP-linked respiration, maximal respiratory capacity, spare capacity, and respiration linked to non-mitochondrial processes were rescued in the presence of CoQ_10_ (Figure 6B-F). Apparent proton leak was reduced in PREPL-deficient cells compared to the control cells (Figure 6G). No significant changes were observed in the treated PREPL-deficient and control cells compared to the non-treated ones.

**Figure 6:**
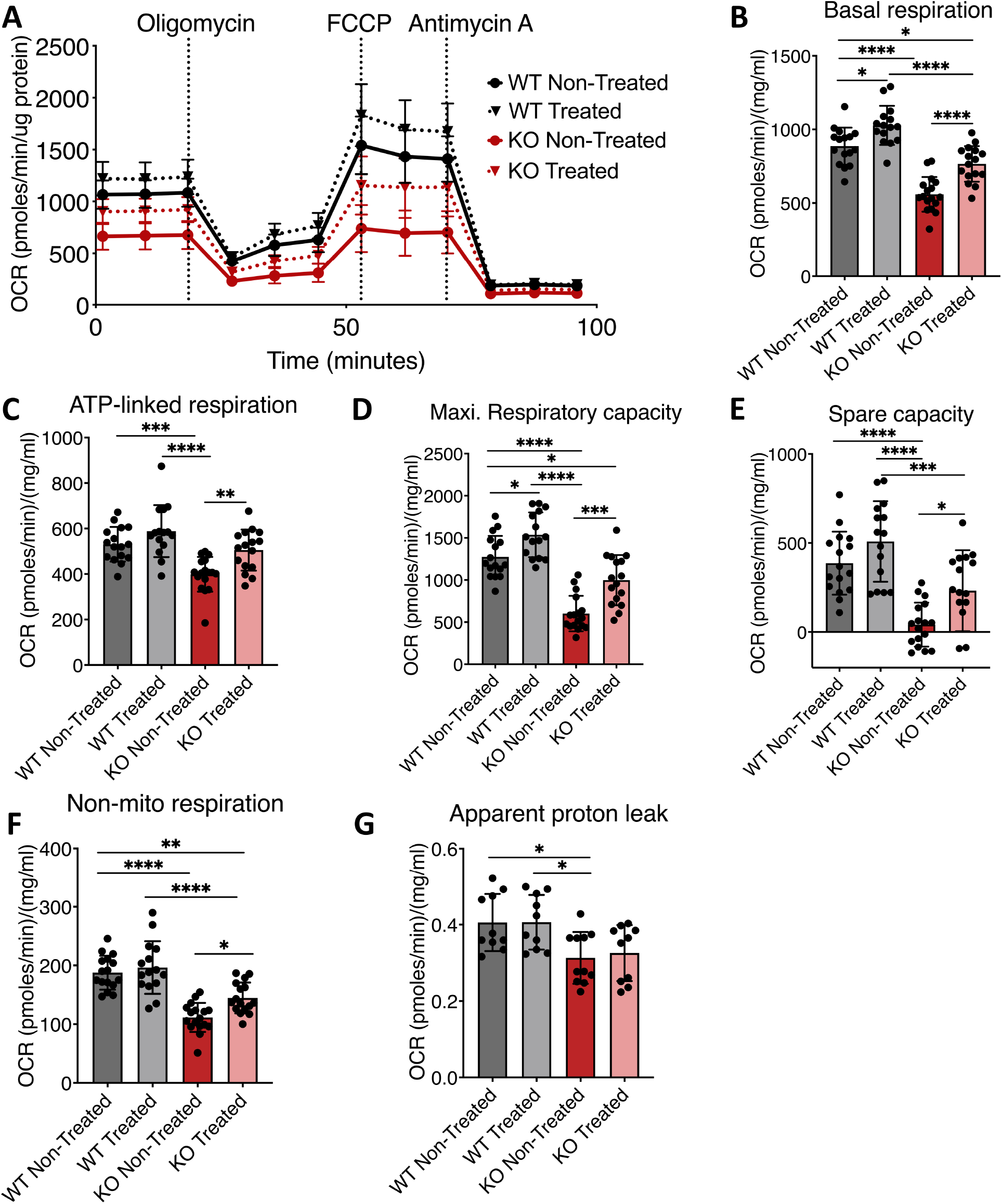
Coenzyme Q_10_ rescues the mitochondrial dysfunction in *PREPL* KO HEK293T cells. (**A**) Respiratory flux profiles indicate that treatment with CoQ (5μM) can stimulate the oxygen consumption rate in *PREPL* KO HEK293T cells compared to the non-treated cells. Bioenergetic parameters (**B**) Basal respiration, (**C**) ATP-linked or coupled respiration (**D**) Maximal respiratory capacity. (**E**) Spare respiratory capacity (**F**) Non-mitochondrial respiration was significantly increased in *PREPL* KO treated cells compared to the non-treated cells. (**G**) Apparent proton leak remained unchanged in treated and non-treated cells. Each data point represents an OCR measurement, plotted as mean ± SD. N > 14 wells for 3 independent experiments. Differences to the controls were analyzed using one-way ANOVA, with Tukey’s test for multiple comparisons. *p < 0.05, **p < 0.01, ***p < 0.001, ****p < 0.0001. All the OCR measurements were normalized to the protein concentration.

### 1.6. PREPL_L_ rescues the mitochondrial dysfunction in PWS patient cells

Lentiviral transduction of the mitochondrial isoform of PREPL (PREPL_L_) was performed in PWS patient skin fibroblast cells to see if this might rescue the mitochondrial dysfunction. Excitingly, the expression of PREPL_L_ rescued the functional capacity of mitochondria in PWS patient cells. This rescue did not appear to be concentration-dependent since a similar rescue was observed for a threefold lower viral transduction. The expression of PREPL in PWS fibroblasts was confirmed using western blot (Supplementary figure 6). OCRs were increased back to wild-type levels in the presence of transduced PREPL_L_ for the bioenergetic parameters basal respiration, ATP-linked respiration, maximal respiratory capacity, and spare capacity (Figure 7A-E). No significant differences were observed in non-mitochondrial respiration amongst all the groups and apparent proton leak was reduced in PWS cells after transduction (Figure 7F-G). No differences were observed in the cells transduced with a control GFP vector, confirming that lentiviral transduction itself has no effect on mitochondrial respiration (Figure 7).

**Figure 7:**
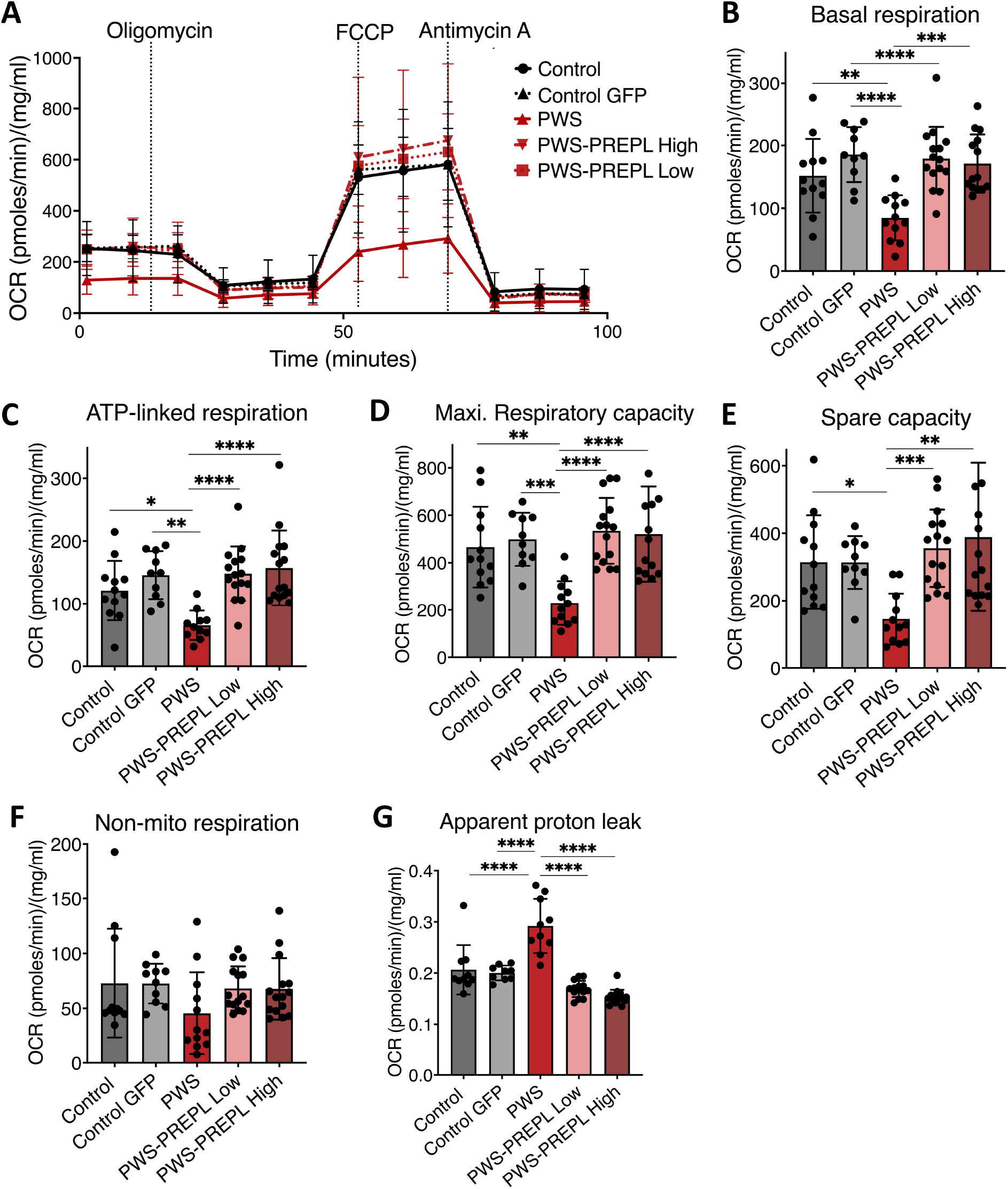
PREPL_L_ rescues mitochondrial dysfunction in PWS patient cells. (**A**) Respiratory flux profiles indicate that expression of PREPL_L_ stimulates oxygen consumption rates in PWS patient fibroblasts. Bioenergetic parameters (**B**) Basal respiration, (**C**) ATP-linked or coupled respiration (**D**) Maximal respiratory capacity (**E**) Spare capacity (**F**) Non-mitochondrial respiration (**G**) Apparent proton leak measured in live cells using mitochondrial stressors. Each data point represents an OCR measurement, plotted as mean ± SD. N > 10 wells for 3 independent experiments. Differences to the controls were analyzed using one-way ANOVA, with Tukey’s test for multiple comparisons. *p < 0.05, **p < 0.01, ***p < 0.001, ****p < 0.0001. All the OCR measurements were normalized to the protein concentration.

## Discussion

In this study, we provide the first molecular links between CMS22 and PWS. Firstly, we have determined that the hypothalamic/pituitary axis is altered in both syndromes, which provides possible explanations for the failure-to thrive phenotype during the neonatal stage and hyperphagia during the adult years. Moreover, expression of *Ghsr1a* was found to be reduced, potentially contributing to short stature and also to impaired cognitive functions (34, 35). Furthermore, mitochondrial bioenergetics are similarly affected in both disorders. We demonstrated that the mitochondrial stimulator Coenzyme Q_10_ can partially rescue the mitochondrial phenotype in PREPL-deficient cells as has previously been suggested for PWS (23, 28). Most excitingly, overexpression of PREPL_L_ rescued the mitochondrial stress and the key metabolic parameters in PWS patient skin fibroblast cells.

*PREPL* expression has been reported to be 2.5-fold downregulated in post-mortem hypothalami of PWS patients (36). Combined with the striking clinical resemblance between CMS22 and PWS and the neonatal hypotonia and impaired growth of *Prepl^-/-^* and PWS-IC^-p/+m^ mice (10, 37), this prompted us to investigate *Prepl* expression in this PWS mouse model. Reduced *Prepl* expression in the neonatal brains of PWS-IC**^-^** ^p/+m^ mice provides a first argument in support of the causal involvement of *Prepl* in PWS phenotype. PWS-IC**^-^**^p/+m^ mice lack expression of all the genes implicated in PWS, and reduced expression of *Prepl* in these mice implies that *Prepl* expression is downstream of another PWS gene. However, the reduction in *Prepl* expression appeared variable, suggesting that the phenotypes in PWS-IC^-p/+m^ mice are caused by PREPL-dependent and -independent mechanisms (see Figure 8 and discussion below). Furthermore, the expression and activity of PREPL protein remain unaltered in peripheral blood lymphocytes of PWS patients (1). The mechanism by which *Prepl* is downregulated is currently unknown but is apparently tissue-specific. The Snord116 cluster has been suggested to contribute to most of the neuroendocrine features observed in PWS patients (38–40). The *in-silico* RNA-RNA interaction prediction algorithm, IntaRNA (41), identified multiple strong interaction motifs between all 30 *SNORD116* copies and *PREPL* mRNA (Supplementary figure 3), suggesting that these can bind and possibly post-transcriptionally stabilize *PREPL* mRNA. Moreover, Baldini et al. identified PREPL as one of the potential targets of Snord116-27 (42). In contrast to the *in-silico* prediction analysis, our study revealed no changes in *Prepl* expression levels in the brain tissue of newborn Snord116^-p/+m^ mice, indicating that PREPL expression is not (solely) regulated by Snord116.

**Figure 8:**
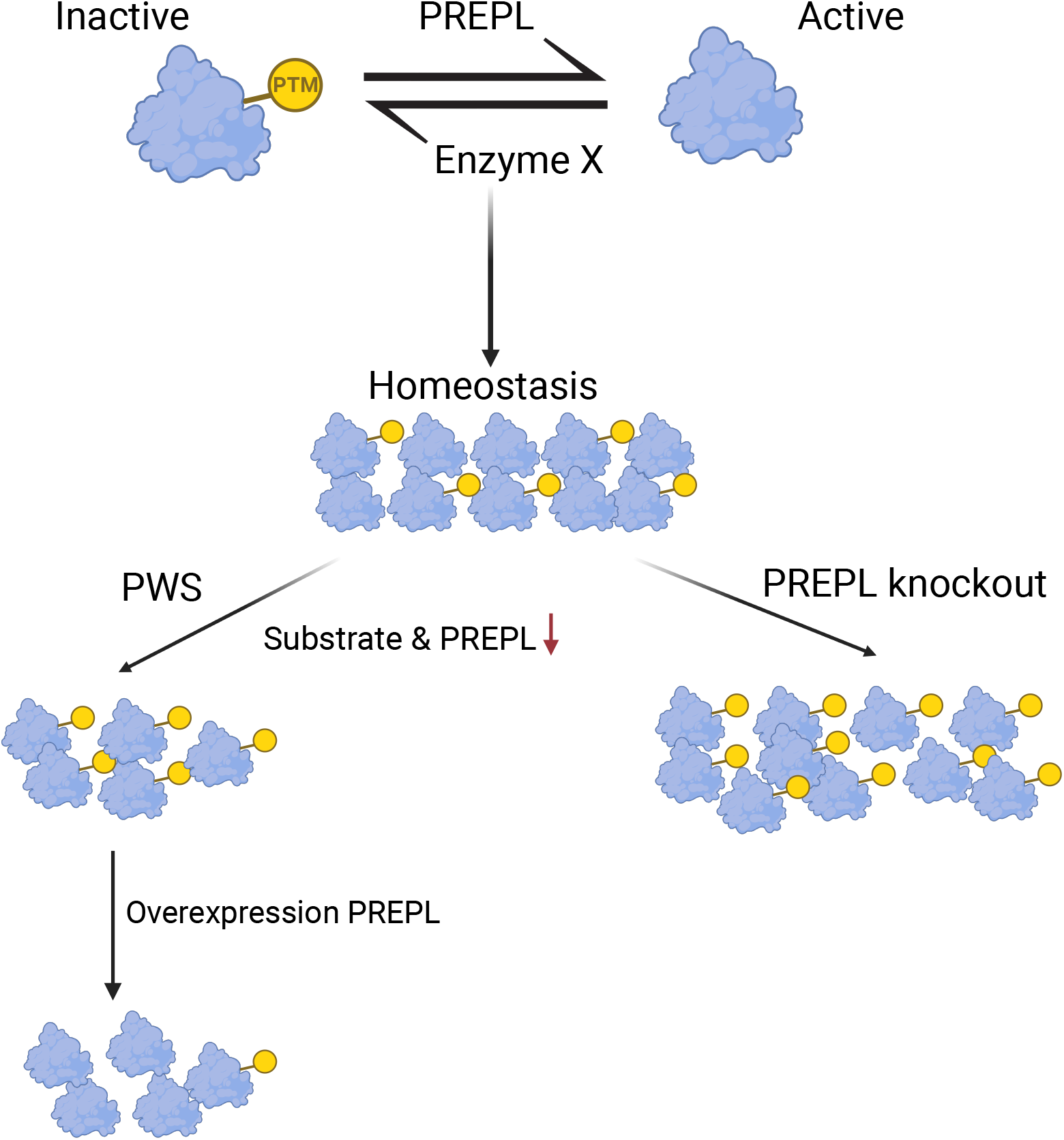
A model of the potential involvement of PREPL in PWS phenotypes. An unknown causal substrate(s) is activated by PREPL by the removal of a post-translational modification (PTM) which is added by an unknown enzyme (indicated as Enzyme X). In CMS22 patients and *Prepl*^-/-^ mice, this leads to reduced amounts of active substrate without affecting the total amount of substrate. In PWS, the amount of active substrate can be reduced in two ways, either through the reduced expression of the substrate or reduced activation due to lower amounts of PREPL. This phenotype can be rescued by PREPL overexpression because the balance is shifted towards active substrate, despite a lower total amount of substrate.

Reduced expression of hypothalamic PC1/3 and one of its transcription factors NHLH2 has been suggested to play a role in several phenotypes of PWS, like reduced short stature, hypogonadism, hyperphagia leading to obesity, hyperghrelinemia and hypothyroidism (4). Since most of these phenotypes are also observed in CMS22, we investigated the expression of *Pcsk1* and *Nhlh2* in both *Prepl*^-/-^ and two different PWS mouse models. We confirmed the downregulation of *Pcsk1* in both *Prepl*^-/-^ and PWS-IC^-p/+m^ brains and *PREPL* KO βTC3 cell lines but not in Snord116^-p/+m^ mice. Since *Nhlh2* shows a trend of upregulation in both the mouse models and is not expressed in the βTC3 cell line, NHLH2 does not appear to be the responsible transcription factor in these mice. Apart from NHLH2, multiple transcription factors including Creb1 and ATF1 are known to transcriptionally regulate the synthesis of *Pcsk1* in hypothalamic/pituitary axis (43–45).

Growth hormone secretagogue receptor 1a (GHSR1a), or ghrelin receptor, regulates hypothalamic-pituitary signalling to stimulate GH release from the pituitary (34). Mutations in GHSR contribute to short stature and delayed pubertal development in patients (35). Moreover, within the arcuate nucleus, this receptor is expressed on NPY/AGRP neurons where it can be activated by acylated ghrelin to inhibit POMC activity, stimulate the expression of *Npy/Agrp* mRNA and subsequently increase food intake (46). Finally, apo-GHSR1a has neuroprotective effects and plays a role in learning and memory (47). Reduced expression of *Ghsr1a* in the pups corresponds well with GH deficiency, short stature, orexigenic defects and intellectual disability observed in these patients. A confounding factor is that reduced expression of the processing enzyme PC1/3 is likely to affect the cleavage of proNPY, AGRP, POMC and other proproteins, which is an essential step in the activation of these precursor proteins (45). Therefore, the net effect on active hormones and neuropeptides remains to be established.

We observed that relative food intake and the expression of the hunger hormone genes *Npy* and *Agrp* were upregulated after refeeding in the adult *Prepl*^-/-^ mice, possibly responsible for hyperphagic behavior and weight gain seen in the patients. Burnett et al. reported similar results during the food intake study performed on adult Snord116^-p/+m^ mice using the same fed/refed conditions (4). We found no indications that the leptin/melanocortin pathway was altered in whole brains of neonatal Snord116^-p/+m^ mice (Supplementary figure 3). Whether this discrepancy is due to age, genetic background, or another factor, is unknown. Collectively, our data suggest that PC1/3 function is impaired in both *Prepl*^-/-^ and PWS-IC^-p/+m^ mouse models and may serve as a contributing factor in PWS and CMS22 phenotypes.

To examine whether PREPL deficiency and PWS are mitochondrial disorders, we analyzed gene expression patterns from the brains of *Prepl*^-/-^ and PWS-IC^-p/+m^ pups using RNAseq. Heatmaps representing statistically significant differentially regulated genes for *Prepl*^-/-^ and PWS-IC^-p/+m^ mice are listed in supplementary figure 4 A and B respectively. Data analysis revealed that many mitochondrial genes are differentially regulated in both disorders, especially those linked to respiratory chain complex machinery and mitoribosomal translation. Defects in a single OXPHOS enzyme can lead to combined deficiencies of respiratory chain supercomplexes as observed in patients with PREPL deficiency and PWS (48). This has also been reported previously in microarray analysis performed on brain tissue of PWS-IC^-p/+m^ mice (24). Our data shows reduced transcription of mtDNA and nuclear DNA encoded complex subunits in the brains of PWS-IC^-p/+m^ mice. In *Prepl*^-/-^ mice respiratory chain complex subunits were increased, but this is most likely a compensatory upregulation since we and others have previously shown that protein levels of these transcripts are reduced due to impaired translation (10).

The mitochondrial function was investigated in PREPL-deficient and PWS cell lines, whereby pharmacological inhibitors of OXPHOS complexes were sequentially added to measure various aspects of respiratory profile (49, 50). In the mitochondrial stress assay, we determined that loss of PREPL in HEK293T cells results in reduced mitochondrial respiration and oxygen consumption rates, thereby affecting OXPHOS complex activities. Additionally, mitochondrial bioenergetics were disturbed in skin fibroblasts of both PWS and PREPL-deficient patients. We observed a significant reduction of basal respiration, ATP-linked respiration, and maximal respiratory capacity, indicating mitochondrial dysfunction. This is in accordance with the previously reported disturbed mitochondrial energy metabolism in PWS and PREPL-deficient patients (23). The number of subjects for this study is small because of the low-throughput nature of the assay. Additional patients with different ages, gender, and ethnicity will be useful in order to validate our findings. On the other hand, the consistency between the HEK293T cell lines, PWS and CMS22 patient fibroblasts is striking and in line with published data (13, 23).

Incubation with CoQ_10_, a potent lipophilic antioxidant rescued mitochondrial dysfunction in PREPL-deficient cells *in vitro*. Supplementation with CoQ_10_ is also known to rescue mitochondrial respiratory chain defects and oxidative stress caused by mtDNA depletion (51, 52). Hence, the involvement of CoQ_10_ in *de novo* pyrimidine synthesis and nucleotide metabolism has the potential to rescue the mitochondrial pathogenicity linked to PWS and CMS22.

Finally, increased expression of PREPL_L_ rescued the altered mitochondrial energy parameters in PWS fibroblast cells. This shows that PWS phenotypes may, at least in part, be directly caused by the downregulation of PREPL. We therefore propose a model for the possible involvement of PREPL in PWS phenotypes (Figure 8). The figure shows that PREPL is either directly responsible for several phenotypes linked to PWS or PREPL is hydrolyzing a post-translational modification on one or more substrates resulting in its activation. Reduced expression of PREPL will lead to reduced activation of these substrates. Alternatively, the substrates are downregulated in PWS but overexpression of PREPL will result in increased hydrolysis and hence activation, leading to a rescue of the phenotype. Given that PREPL cleaves ester and thioester substrates (10), potential substrates are palmitoylated and acetylated proteins.

In conclusion, this study identifies two novel molecular links between CMS22 and PWS. Impaired hypothalamic/pituitary signaling is responsible for the neuroendocrine phenotypes in both syndromes. In addition, impaired mitochondrial bioenergetics and respiratory chain defects suggest that mitochondrial dysfunction contributes to the clinical phenotype of both disorders. Moreover, the mitochondrial deficiency in PWS cells can be rescued by PREPL_L_. Given the molecular overlap between the disorders, the identification of physiological substrates of PREPL might lead to novel therapeutic targets for both syndromes.

## Material and methods

### Animal models (Mouse breeding and genotyping)

#### 1. Breeding

*Prepl* KO (*Prepl*^-/-^) mice were generated by targeted deletion of exon 10 of *Prepl* using homologous recombination in embryonic stem cells (ES), as described previously (10). These mixed background (50% C57BL/6, 25%129S, 25%SW) mice were obtained by backcrossing (Prepl-ex10(Fl) tm Lox KO) to C57BL/6 once to avoid the neonatal lethality observed in pure C57BL/6 background.

PWS^-p/+m^ mice (C57BL/6J-35KBdelPWS-IC) have also been described before (53). This PWS mouse model was generated by standard ES cell targeting and shows similar defects in paternally imprinted genes as observed in PWS. Since the mutation results in postnatal lethality when inherited from males on C57BL/6J background (30),the line was maintained by crossing IC^-p/+m^ females with WT males.

Floxed Snord116^-p/+m^ mice (B6.Cg-Snord116tm1Uta/J, Jax stock 008118) (54) were obtained from the University of Manchester. Female offspring were crossed to CMV-cre mice (B6.C-Tg(CMV-cre)1Cgn/J, Jax stock 006054) (55) and a female carrying a deleted Snord116 allele was crossed to cmv-cre mice one more time. The resulting mice were crossed to wildtype C57BL/6N (Charles River) 10 times to backcross to a clean genetic background and to lose potential unrecombined alleles. Males carrying the deleted allele were then mated with female wild-type C57BL/6N mice. The male breeder was removed from the breeding cage after 10 days.

The mice were housed under SPF conditions in standard cages on a 12h light/darkness cycle, with *ad libitum* access to water and a diet of standard rodent chow. The brains were isolated from *Prepl*^-/-^, PWS^-p/+m,^ and Snord116^-p/+m^ pups within 4-5 hours after birth, snap-frozen in liquid nitrogen, and stored at −80°C. All the studies were done using littermate controls.

#### 2. Genotyping

Genotyping of the *Prepl*^-/-^ and PWS^-p/+m^ mice was carried out on tail/ ear genomic DNA (10). For Snord116^-p/+m^ mice, a tail biopsy taken at sacrifice on the day of birth was cooked in 50 mM NaOH at 98 °C for 1h, followed by the addition of 50ul 1M Tris-HCl. 1.25µl was used for the PCR reaction employing GoTaq G2 DNA Polymerase (M7845, Promega, Dübendorf, Switzerland) with primers (Microsynth, Balgach, Switzerland) on a Biometra TRIO thermocycler (Bartelt GmbH, Innsbruck, Austria) and visualized on agarose gels using electrophoresis. Primers used for mouse genotyping are specified in Supplementary table 1.

### Cell culture and patient skin fibroblasts

HEK293T and βTC3 cells were obtained from the American Type Culture Collection (ATCC, Manassas, VA, USA). Human skin fibroblast cells were established from two patients with PREPL deficiency (males, aged between 2 weeks and 7 months) (56), two Prader-Willi Syndrome with deletion (males, aged between 10-42 years), and two healthy controls for the mitochondrial functional studies (57).

Fibroblast cell cultures of patients and controls were established as follows: Skin biopsies were cut into small pieces and transferred to sterile 25 cm^2^ tissue culture flasks (Falcon). 4 ml AmnioPAN culture medium (Pan-Biotech) without serum was added followed by the addition of 1 ml collagenase (2000 units/ml (Pan-Biotech)). The cultures were then maintained overnight at 37°C in a CO_2_ incubator. Next day, cells were trypsinized and centrifuged for 10 minutes at 122g, 4°C. The pellet was resuspended in 10 ml AmnioPAN medium without serum in 25 cm^2^ flasks and maintained at 37°C and 5% CO_2_. The medium was refreshed twice a week.

The cells were grown in DMEM (Thermo Fisher Scientific) supplemented with HEPES and 10% fetal calf serum (FCS) (VWR) 37°C/5% CO2. Cells were used for experiments between passages 4–7 and maintained at ∼75-80% confluency at 37°C, 5% CO_2_ in a humidified atmosphere.

For rescue experiments, HEK293T cells were incubated with 5 μM of Coenzyme Q_10_ (CoQ_10_; Pharma Nord) for 5 days and then plated for mitochondrial assays. CoQ_10_ was solubilized in ethanol and was diluted 2000-fold in growth media for at least 15 minutes at 37°C to ensure adequate absorption (26).

### CRISPR-Cas9

*PREPL* knockout (KO) HEK293T and βTC3 cell lines were generated with the CRISPR-Cas9 genome engineering system following the protocol described by Ran et al. (58–60). Briefly, for HEK293T cells, a guide sequence targeting exon 4 and 5 was cloned in the pSpCas9(BB)-2A-Puro construct (Addgene 48139). For βTC3 cells, a guide sequence targeting *Prepl* exon 3 and intron 3 was cloned in the same construct. After the transfection, puromycin selection was performed on cells for three days. Cells were serially diluted in 96-well plates to obtain single cells. To check the efficiency of the guide, genomic DNA was extracted from transfected cells using the QuickExtract^TM^ DNA Extraction Solution (Lucigen) and mutations were detected by the SURVEYOR assay (Integrated DNA technologies). Genomic aberrations were verified by TOPO cloning followed by Sanger sequencing. PCR product was purified using the QIAquick PCR purification kit (Qiagen) followed by A-tailing and ligation into a pGEM®-T Easy vector. PREPL deficiency in all the clones was verified by Western blot using anti-PREPL antibody (Santa Cruz). Primers for cloning of the guide sequence and Sanger sequencing for HEK293T and βTC3 cells are listed in Supplementary table 2 and 3 respectively.

### Lentiviral transduction

A 2.2 kb BamHI-SalI cDNA fragment of the long form of human *PREPL* (*PREPL_L_*) was cloned into pLenti-GFP (Addgene) digested with the same restriction enzymes, which replaced the GFP cDNA present in the vector. To produce lentiviral vectors overexpressing PREPL_L_ or GFP, packaging was performed in HEK293T cells following the standard protocols. HEK293T cells were transfected with three different plasmids: pLenti-*PREPL_L_* or pLenti-*GFP*, together with pVSVG, encoding envelop protein: vesicular stomatitis G protein (VSVG) and pdelta 8.91, encoding packaging protein delta 8.91. Lentiviral media was harvested and filtered using 0.45 nm filter 48 hours post-transfection and centrifuged for 5 minutes at maximum speed. The lentiviral media was diluted in complete medium to reach 1:3 (high) and 1:10 (low) concentrations. PWS and control skin fibroblasts were transduced with lentiviral media and polybrene (8 µg/ml) (1:1 ratio of viral particle to polybrene medium). The cells were replaced with the complete medium after 4 hours. After 3 days, cells were selected with puromycin (1 µg/ml) for another 5 days to generate stable cell lines.

### Real-time quantitative PCR (RT-qPCR)

RNA was isolated from brain of neonatal *Prepl*^-/-^ and PWS^-p/+m^ pups and their wildtype littermates using the Nucleospin RNA II kit (Macherey Nagel), from hypothalamus using the Nucleospin RNA XS kit (Macherey Nagel). The quality of the RNA was assessed using a NanoDrop ND-1000 spectrophotometer (V.3.8.1). cDNA was synthesized with the iScript cDNA synthesis kit (BioRad). Real-time PCR was performed in triplicate with the CFX Connect^TM^ Real-Time PCR Detection system (BioRad) using SsoAdvanced^TM^ Universal SYBR® Green Supermix (BioRad). Data were analyzed using the Livak method (61). Samples were normalized to glyceraldehyde 3-phosphate dehydrogenase (*Gapdh*). Primers used for RT-qPCR are listed in Supplementary table 4.

### Food intake analysis and isolation of hypothalami

For food intake experiments, *Prepl*^-/-^ mice were transferred to grid cages 3 days before starting food intake measurements for acclimatization. For 5 subsequent days, the food eaten and spilled was measured every 24 hours. Mice were sacrificed by cervical dislocation after 16h overnight (O/N) fasting or O/N fasted + refed for 5 hours before the hypothalamus was isolated. Hypothalami were snap-frozen in liquid nitrogen and stored at −80°C.

### Mitochondrial bioenergetics

Mitochondrial bioenergetics were analyzed by measuring oxygen consumption rates (OCR) using the Agilent XF24 extracellular flux analyzer (Seahorse Bioscience, North Billerica, MA) according to the guidelines of the manufacturer (62). Briefly, HEK293T and skin fibroblast cells were seeded in XF24 cell culture plates at a density of 45,000 cells/well and 30,000 cells/well, respectively, in DMEM containing 10% FCS). After 20- 24 hours incubation, adherent cells were washed twice with pre-warmed XF base medium supplemented with 10 mM glucose (HEK293T) or 25mM glucose (skin fibroblasts), 2 mM glutamine, and 1 mM sodium pyruvate; pH 7.4) and incubated for 1h at 37 °C. The sensor cartridge was hydrated overnight at 37°C with XF Calibrant and calibrated using the XF24 analyzer. Oxygen consumption rates (OCRs) were measured under baseline conditions followed by the sequential addition of oligomycin, carbonyl cyanide trifluoromethoxyphenylhydrazone (FCCP) and antimycin A to injector ports A, B, and C of the cartridge, respectively. The final concentrations of the injections were 2 μM oligomycin, 0.5 μM FCCP, and 1 μM antimycin A for HEK293T and 10 μM oligomycin, 2.7 μM FCCP, and 10 μM antimycin A for skin fibroblast cells. OCRs were normalized to the protein concentrations determined by the Pierce BCA Protein assay kit (Thermo Fisher Scientific).

### RNA sequencing

RNA from neonatal brains of *Prepl*^-/-^ (n=4), PWS^-p/+m^ (n=4) and their respective control littermates (n=4) was sequenced. The RNA integrity number (RIN-value; degradation parameter) was determined using the Agilent 2100 Bioanalyzer (Agilent Technologies, Inc., Santa Clara, CA, USA) and had values ≥ 7. 500ng of high-quality input sample was used to initiate cDNA-cRNA synthesis via reverse transcription using Quantseq 3 mRNA library prep kit (Lexogen). RNA was removed using removal solution-Globin block (RS-GBHs) and the library was amplified using single indexing (i7 only) PCR. Libraries were sequenced on Illumina HiSeq4000 with 1 x 50 bp read length and a minimum 2 million reads per sample. Reads were mapped to reference genome (house mouse; NCBI/build 38). Quality control of raw reads was performed with FastQC v0.11.7 (63) and adapters were filtered with ea-utils fastq-mcf v1.05. Splice-aware alignment was performed with HISAT2 (64) against the reference genome mus musculus 10 using the default parameters. Reads mapping to multiple loci in the reference genome were discarded. The resulting binary alignment map files (BAM) were handled with Samtools v1.5 (65). Quantification of reads per gene was performed using HT-seq Count v0.10.0, Python v2.7.14 (66). The standard count-based differential expression analysis was done with R-based (The R Foundation for Statistical Computing, Vienna, Austria) Bioconductor package DESeq2 (67). Reported p-values were adjusted for multiple testing with the Benjamini-Hochberg procedure, which controls the false discovery rate (FDR). The gene expression profiles were analyzed using Ingenuity Pathway Analysis (IPA) software. The top 50 genes were selected based on the adjusted p-value for differential expression. Only mitochondrial genes with adjusted p-value <0.1 were considered. Heatmaps were created with Heatmapper, scaling each gene, with average linkage clustering based on Euclidean distance measurement (68).

### Western Blot

Snap-frozen brains from *Prepl*^-/-^, PWS-IC^-p/+m^, Snord116^-p/+m^, and control pups were homogenized by sonication in Cell Lysis Buffer (Cell Signaling Technology) supplemented with complete EDTA-free protease inhibitors (Roche). HEK293T and βTC3 cells were homogenized in RIPA buffer supplemented with complete EDTA-free protease inhibitors. Proteins were separated by SDS-PAGE and analyzed by Western blotting using anti-PREPL (Invitrogen), anti-PC1/3 (homemade), and anti-GAPDH (Cell Signaling Technology) antibodies. Proteins were detected with the Western Lightning ECL system (Perkin Elmer) and images were quantified using ImageJ software (69).

### Quantification and Statistical Analysis

All statistical analyses were performed with GraphPad Prism 9. Statistical analysis was performed by unpaired two-sided Student’s t-tests for two groups and one-way ANOVA with Tukey’s multiple comparisons for more than two groups. Data are represented as mean ± SD. Significance shown on graphs as *p < 0.05, **p < 0.01 or ***p < 0.001. Statistical details can be found in the figure legends.

### Study approval

All the animal studies were approved by the ethical research committee of KU Leuven in accordance with the declaration of Helsinki (Project number, 282/2015), Institutional Animal Care and Use Committee (IACUC), University of Florida (Project number, 202210656) and by the institutional authorities of Basel in accordance with the federal laws of Switzerland (Project number, 35195/3045). For taking the skin fibroblast samples from patients, written consent was received prior to participation.

## Author contributions

KB, KR, YM and SM conducted the biochemical experiments. KB performed and analyzed mitochondrial stress studies supervised by EV and PA. LT analyzed the RNAseq data and generated the heat maps. JWC and KR designed and characterized the *Prepl*^-/-^ mouse model. JLR designed, characterized, and provided the tissues for PWS-IC^-p/+m^ mouse model. DTM designed, characterized, and performed RT-qPCR studies on Snord116^-p/+m^ mouse model. JWC and AR designed the research. KB and JWC collected and analyzed the data and wrote the manuscript with inputs from all the authors.

## Acknowledgments

This work was supported by ‘Stichting Marguerite-Marie Delacroix’ and ‘FWO Vlaanderen’ (#GOB9119N). We would like to thank KU Leuven C14/21/095 InterAction consortium for the purchase of Seahorse Analyzer used for mitochondrial studies.

## References

1. Régal L, et al. PREPL deficiency: delineation of the phenotype and development of a functional blood assay. Genetics in Medicine. 2018;20(1):109–118.

2. Duarte ML, et al. Multiomics Analyses Identify Proline Endopeptidase-Like Protein As a Key Regulator of Protein Trafficking, a Pathway Underlying Alzheimer’s Disease Pathogenesis S. 10.1124/molpharm.122.000641.

3. Juriaans AF, Kerkhof GF, Hokken-Koelega ACS. The Spectrum of the Prader– Willi-like Pheno- and Genotype: A Review of the Literature. Endocr Rev. 2022;43(1):1–18.

4. Burnett LC, et al. Deficiency in prohormone convertase PC1 impairs prohormone processing in Prader-Willi syndrome. J Clin Invest. 2017;127(1):293–305.

5. Polex-Wolf J, et al. Hypothalamic loss of Snord116 recapitulates the hyperphagia of Prader-Willi syndrome. J Clin Invest. 2018;128(3):960.

6. Seidah NG. Proprotein Convertase 1/3. Handbook of Proteolytic Enzymes. 2013;3:3286–3290.

7. Thorner E. Disruption of PC13 expression in mice causes dwarfism and multiple neuroendocrine peptide processing defects. 1999.

8. Ramos-Molina B, et al. Hyperphagia and obesity in prader–willi syndrome: PCSK1 deficiency and beyond? Genes (Basel). 2018;9(6). 10.3390/genes9060288.

9. Jackson, Robert S.; Creemers JWM;, et al. Obesity and impaired prohormone processing associated with mutations in the human prohormone convertase 1 gene. Nat Genet. 1997;16(July):303–306.

10. Rosier K, et al. Prolyl endopeptidase-like is a (thio)esterase involved in mitochondrial respiratory chain function. iScience. 2021;24(12). 10.1016/J.ISCI.2021.103460.

11. Morawski M, et al. Cellular and ultra structural evidence for cytoskeletal localization of prolyl endopeptidase-like protein in neurons. Neuroscience. 2013;242:128–139.

12. Rosier K, Creemers J. Unraveling the role of PREPL, a serine hydrolase deleted in a Prader-Willi-like syndrome [Doctoral dissertation, KU Leuven]. 2021.

13. Kramer NJ, et al. Regulators of mitonuclear balance link mitochondrial metabolism to mtDNA expression. Nat Cell Biol. [published online ahead of print: 2023]. 10.1038/S41556-023-01244-3.

14. Bartholdi D, et al. Further delineation of genotype-phenotype correlation in homozygous 2p21 deletion syndromes: first description of patients without cystinuria. Am J Med Genet. 2013;161A(8):1853–9.

15. Legati A, et al. New genes and pathomechanisms in mitochondrial disorders unraveled by NGS technologies. Biochim Biophys Acta Bioenerg. 2016;1857(8):1326–1335.

16. Martens K, et al. Global distribution of the most prevalent deletions causing hypotonia–cystinuria syndrome. European Journal of Human Genetics. 2007;15(10):1029–1033.

17. Wortmann SB, et al. Whole exome sequencing of suspected mitochondrial patients in clinical practice. J Inherit Metab Dis. 2015;38(3):437–443.

18. Ng YS, Turnbull DM. Mitochondrial disease: genetics and management. J Neurol. 2016;263(1):179.

19. Khan NA, et al. Mitochondrial disorders: challenges in diagnosis & treatment. Indian J Med Res. 2015;141(1):13–26.

20. Perrone S, et al. Oxidative Stress in Cancer-Prone Genetic Diseases in Pediatric Age: The Role of Mitochondrial Dysfunction. [published online ahead of print: 2016]. 10.1155/2016/4782426.

21. Filiano JJ, et al. Mitochondrial Dysfunction in Patients With Hypotonia, Epilepsy, Autism, and Developmental Delay: HEADD Syndrome. J Child Neurol. 2002;17:435– 439.

22. Towheed A, et al. Hypotonia–cystinuria 2p21 deletion syndrome: Intrafamilial variability of clinical expression. Ann Clin Transl Neurol. 2021;8(11):2199–2204.

23. Butler MG, et al. Preliminary Observations of Mitochondrial Dysfunction in Prader-Willi Syndrome. Am J Med Genet A. 2018;176(12):2587.

24. Yazdi PG, et al. Differential gene expression reveals mitochondrial dysfunction in an imprinting center deletion mouse model of prader-willi syndrome. Clin Transl Sci. 2013;6(5):347–355.

25. Hargreaves IP. Coenzyme Q10 as a therapy for mitochondrial disease. International Journal of Biochemistry and Cell Biology. 2014;49(1):105–111.

26. Duberley KE, et al. Effect of Coenzyme Q10 supplementation on mitochondrial electron transport chain activity and mitochondrial oxidative stress in Coenzyme Q 10 deficient human neuronal cells. International Journal of Biochemistry and Cell Biology. 2014;50(1):60–63.

27. Butler MG, et al. Coenzyme Q10 Levels in Prader-Willi Syndrome: Comparison With Obese and Non-Obese Subjects. Am J Med Genet A. 2003;119A(2):168–171.

28. Eiholzer U, et al. Developmental profiles in young children with Prader-Labhart-Willi syndrome: Effects of weight and therapy with growth hormone or coenzyme Q 10. Am J Med Genet A. 2008;146(7):873–880.

29. Yang T, et al. A mouse model for Prader-Willi syndrome imprinting-centre mutations. 1998.

30. Chamberlain SJ, et al. Evidence for genetic modifiers of postnatal lethality in PWS-IC deletion mice. 10.1093/hmg/ddh314.

31. Adhikari A, et al. Cognitive deficits in the Snord116 deletion mouse model for Prader-Willi syndrome. Neurobiol Learn Mem. 2019;165(April 2018):106874.

32. Kann O, Kovács R. Mitochondria and neuronal activity. Am J Physiol Cell Physiol. 2007;292(2):641–657.

33. Top-Down Control Analysis of Systems with More than one Common Intermediate [Internet].

34. Yin Y, Li Y, Zhang W. The growth hormone secretagogue receptor: Its intracellular signaling and regulation [preprint]. Int J Mol Sci. 2014;15(3):4837–4855.

35. Barroso PS, et al. Clinical and Genetic Characterization of a Constitutional Delay of Growth and Puberty Cohort. Neuroendocrinology. 2020;110(11–12):959–966.

36. Falaleeva M, et al. SNORD116 and SNORD115 change expression of multiple genes and modify each other’s activity. Gene. 2015;572(2):266–273.

37. Lone AM, et al. Deletion of Prepl Causes Growth Impairment and Hypotonia in Mice. PLoS One. 2014;9(2):89160.

38. Qi Y, et al. Snord116 is critical in the regulation of food intake and body weight. Sci Rep. 2016;6. 10.1038/SREP18614.

39. Duker AL, et al. Paternally inherited microdeletion at 15q11.2 confirms a significant role for the SNORD116 C/D box snoRNA cluster in Prader–Willi syndrome. European Journal of Human Genetics. 2010;18:1196–1201.

40. Grootjen LN, et al. Atypical 15q11.2-q13 Deletions and the Prader-Willi Phenotype. J Clin Med. 2022;11(15). 10.3390/jcm11154636.

41. Mann M, Wright PR, Backofen R. IntaRNA 2.0: enhanced and customizable prediction of RNA–RNA interactions. Nucleic Acids Res. 2017;45(Web Server issue):W435.

42. Baldini L, et al. Phylogenetic and Molecular Analyses Identify SNORD116 Targets Involved in the Prader-Willi Syndrome. 10.1093/molbev/msab348.

43. Greenwood M, et al. Transcription factor Creb3l1 regulates the synthesis of prohormone convertase enzyme PC1/3 in endocrine cells. J Neuroendocrinol. 2020;32. 10.1111/jne.12851.

44. Jansen E, et al. Cell Type-specific Protein-DNA Interactions at the cAMP Response Elements of the Prohormone Convertase 1 Promoter. Journal of Biological Chemistry. 1997;272(4):2500–2508.

45. Stijnen P, et al. PCSK1 Mutations and Human Endocrinopathies: From Obesity to Gastrointestinal Disorders. [published online ahead of print: 2016]. 10.1210/er.2015-1117.

46. Briggs DI, et al. Diet-Induced Obesity Causes Ghrelin Resistance in Arcuate NPY/AgRP Neurons. [published online ahead of print: 2010]. 10.1210/en.2010-0556.

47. Kern A, Grande C, Smith RG. Apo-Ghrelin Receptor (apo-GHSR1a) Regulates Dopamine Signaling in the Brain. Front Endocrinol (Lausanne). 2014;5(AUG). 10.3389/FENDO.2014.00129.

48. Mayr JA, et al. Spectrum of combined respiratory chain defects. J Inherit Metab Dis. 2015;38(4):629–640.

49. Brand MD, Nicholls DG. Assessing mitochondrial dysfunction in cells. Biochem J. 2011;435:297–312.

50. Dranka BP, et al. Assessing bioenergetic function in response to oxidative stress by metabolic profiling. [published online ahead of print: 2010]. 10.1016/j.freeradbiomed.2011.08.005.

51. López-Martín JM, et al. Missense mutation of the COQ2 gene causes defects of bioenergetics and de novo pyrimidine synthesis. Hum Mol Genet. 2007;16(9):1091– 1097.

52. Duberley KE, et al. Effect of Coenzyme Q10 supplementation on mitochondrial electron transport chain activity and mitochondrial oxidative stress in Coenzyme Q 10 deficient human neuronal cells. International Journal of Biochemistry and Cell Biology. 2014;50(1):60–63.

53. Yang T, et al. A mouse model for Prader-Willi syndrome imprinting-centre mutations. 1998.

54. Ding F, et al. SnoRNA Snord116 (Pwcr1/MBll-85) deletion causes growth deficiency and hyperphagia in mice. PLoS One. 2008;3(3). 10.1371/journal.pone.0001709.

55. Schwenk F, Baron U, Rajewsky K. A cre-transgenic mouse strain for the ubiquitous deletion of loxP-flanked gene segments including deletion in germ cells. Nucleic Acids Res. 1995;23(24):5080–5081.

56. Jaeken J, et al. Deletion of PREPL, a gene encoding a putative serine oligopeptidase, in patients with hypotonia-cystinuria syndrome. Am J Hum Genet. 2006;78(1):38–51.

57. Dranka BP, et al. Assessing bioenergetic function in response to oxidative stress by metabolic profiling. [published online ahead of print: 2010]. 10.1016/j.freeradbiomed.2011.08.005.

58. Rosier K, Creemers J. Unraveling the role of PREPL, a serine hydrolase deleted in a Prader-Willi-like syndrome [Doctoral dissertation, KU Leuven]. 2021.

59. Monnens Y, Rosier K, Creemers J. Functional characterization of PREPL variants : implications of missense mutations on catalytic activity, protein-protein interactions and cellular functioning [Master’s thesis, KU Leuven]. 2021.

60. Ran FA, et al. Genome engineering using the CRISPR-Cas9 system. Nat Protoc. 2013;8(11):2281–2308.

61. Livak KJ, Schmittgen TD. Analysis of relative gene expression data using real-time quantitative PCR and the 2(-Delta Delta C(T)) Method. Methods. 2001;25(4):402–408.

62. Nicholls DG, et al. Bioenergetic Profile Experiment using C2C12 Myoblast Cells. JoVE. 2010;46. 10.3791/2511.

63. S. A. FASTQC. A quality control tool for high throughput sequence data. 2010. [Internet].

64. Kim D, et al. Graph-Based Genome Alignment and Genotyping with HISAT2 and HISAT-genotype. Nat Biotechnol. 2019;37(8):907.

65. Li H, et al. The Sequence Alignment/Map format and SAMtools. BIOINFORMATICS APPLICATIONS NOTE. 2009;25(16):2078–2079.

66. Anders S, Pyl PT, Huber W. Genome analysis HTSeq-a Python framework to work with high-throughput sequencing data. 2015;31(2):166–169.

67. Love MI, Huber W, Anders S. Moderated estimation of fold change and dispersion for RNA-seq data with DESeq2. Genome Biol. 2014;15:550.

68. Babicki S, et al. Heatmapper: web-enabled heat mapping for all. Nucleic Acids Res. 2016;44:147–153.

69. Schneider CA, Rasband WS, Eliceiri KW. NIH Image to ImageJ: 25 years of Image Analysis. Nat Methods. 2012;9(7):671.

