## Supplementary data for "Similar metabolic pathways are affected in both Congenital Myasthenic Syndrome-22 and Prader-Willi Syndrome"

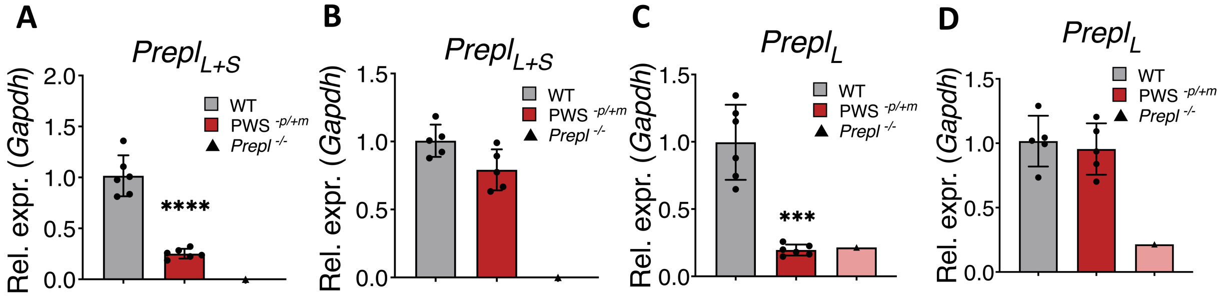


**Supplementary figure 1:** mRNA levels of *Prepl* in two different cohorts of PWS-IC$^{-p/+m}$ mice. (**A, C**) Reduced gene expression of *Prepl*$_{L+S}$ and *Prepl*$_{L}$ in the first cohort, and (**B, D**) Unaltered gene expression of *Prepl*$_{L+S}$ and *Prepl*$_{L}$ in the second cohort. Each data point represents 1 animal (N > 4 per genotype, N = 1 *Prepl*$^{-/-}$); plotted as mean ± SD. Differences to the unaffected controls were analyzed with one-way ANOVA with Tukey's test for multiple comparisons. ***$p < 0.001$, ****$p < 0.0001$.

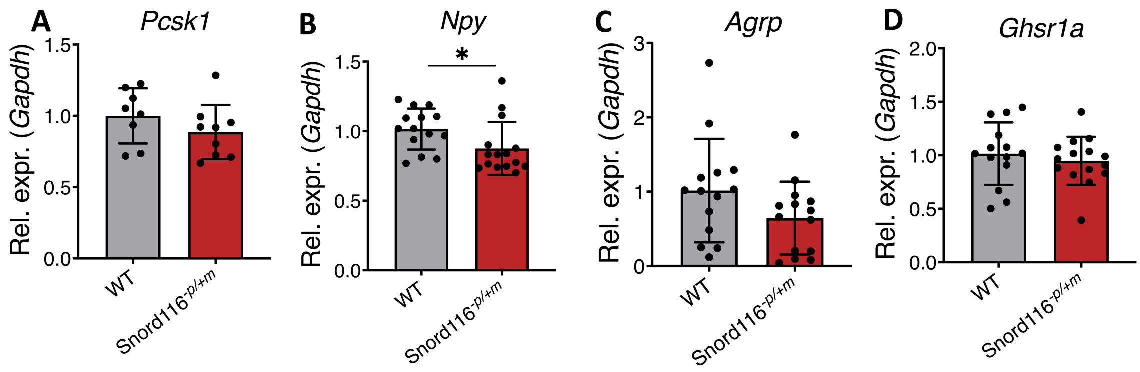


**Supplementary figure 2: Hypothalamic/ pituitary axis remains unaffected in Snord116$^{-p/+m}$ model.** (**A**) Transcript levels of *Pcsk1* in the whole brains of Snord116$^{-p/+m}$ pups. (**B** and **C**) mRNA levels of *Npy* are reduced while *Agrp* levels are not significantly affected in Snord116$^{-p/+m}$ mice. (**D**) Gene expression levels of *Ghsr1a* in neonatal Snord116$^{-p/+m}$ mice. Each data point represents 1 animal (N > 6 per genotype), plotted as mean ± SD. Differences to the unaffected controls were analyzed with a 2-tailed, type 3 (assumes unequal variance) Student's t test. *$p < 0.05$.

**A**

```
                          983                                                             1018
                           |                                                               |
PREPL 3' UTR   5'-AAC..AAAAU        A           UAAA        U                AUG          GCUCU..AUU-3'
                           UCAGA   CCA  CGA      GAUUC             GUCG      CCAUCCA
                           |||||   |||  |||      |||||             ||||      |||||||
                           AGUCU   GGU  GCU      CUAAG             CAGC      GGUAGGU
SNORD116-28    3'-AAC..AAACG       AA     U             UAAAAAUU        AGUA
                           |                                                               |
                          42                                                               1
```

**B**

| | *PREPL 3' UTR interaction energy (kcal/mol)* | | | *PREPL mRNA interaction energy (kcal/mol)* | | |
|---|---|---|---|---|---|---|
| ***SNORD116-23*** | **-10,89** | | | **-11,11** | **-10,6** | **-10,25** |
| ***SNORD116-28*** | **-10,91** | **-10,67** | **-9,7** | **-10,77** | | |

**Supplementary figure 3: Strong interaction motifs exist between *SNORD116* and *PREPL*.** (**A**) Predicted interaction strengths of multiple predicted interaction domains between *SNORD116-23, SNORD116-28* and *PREPL*. (**B**) Visual representation of the strongest interaction domain between *SNORD116-28* and the *PREPL* 3'UTR.

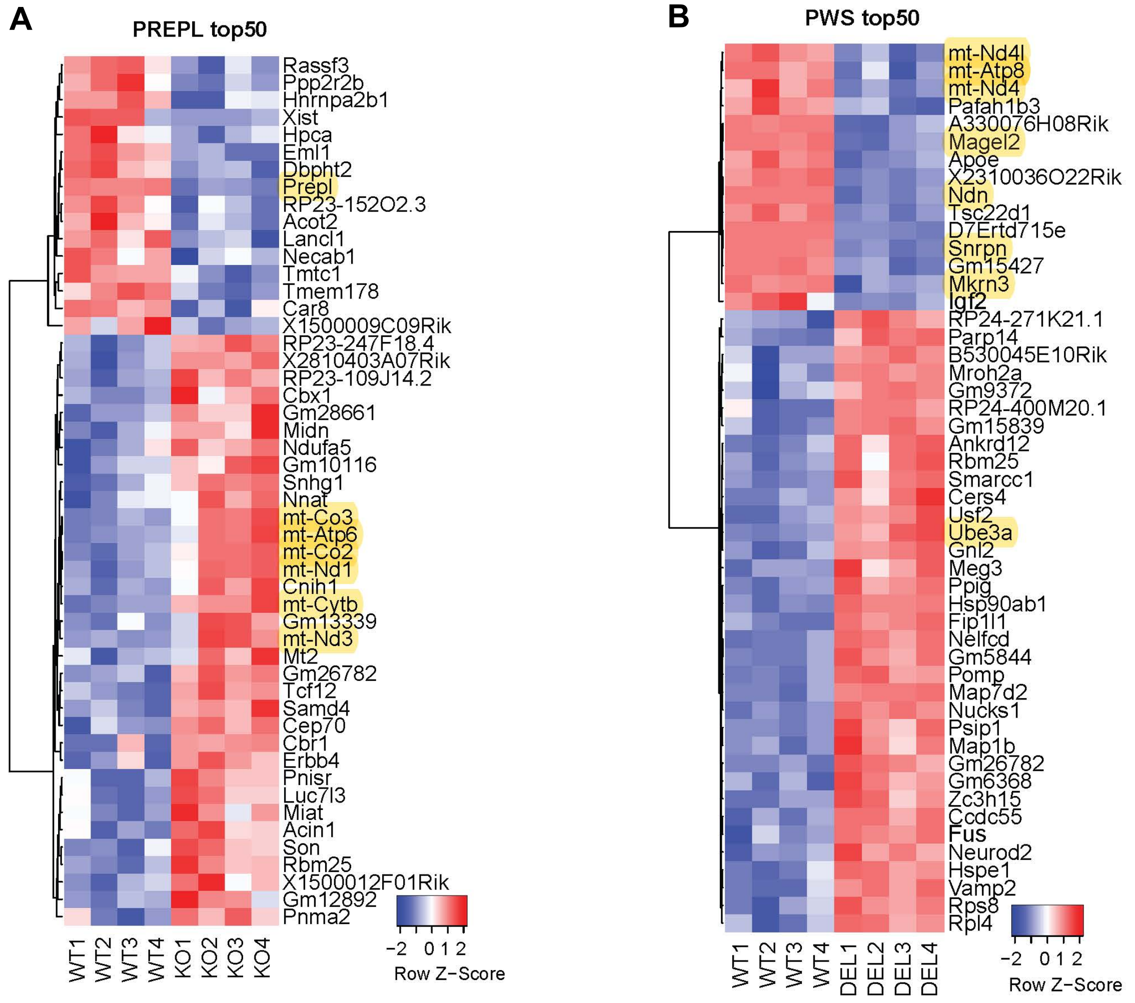


**Supplementary figure 4:** Heatmap representing top 50 differentially expressed genes in the brains of *Prepl*$^{-/-}$ and PWS-IC$^{-p/+m}$ mice.

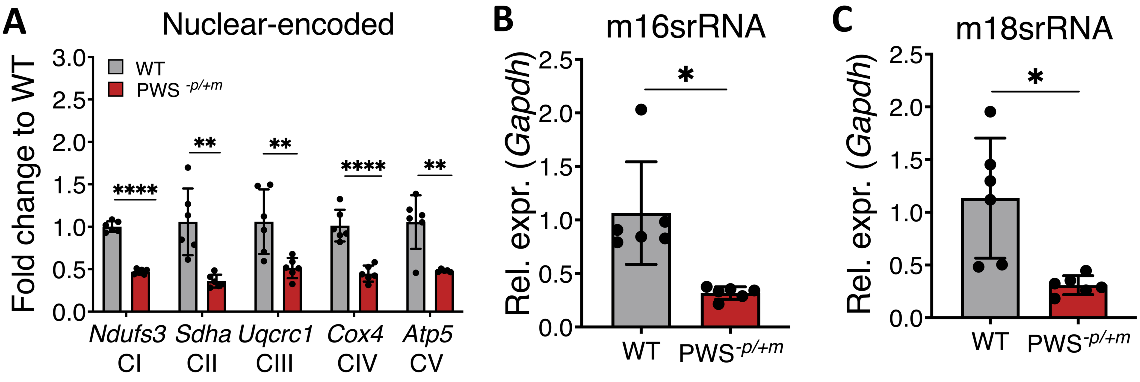


**Supplementary figure 5:** Gene expression levels of mitochondrial-linked genes in PWS-IC$^{-p/+m}$ mice. (**A**) Decreased transcription of nuclear-encoded complex subunits in PWS-IC$^{-p/+m}$ mice. (**B**) Quantification of mitochondrially encoded 16s rRNA in PWS-IC$^{-p/+m}$ mice. (**C**) Quantification of mitochondrially encoded 18s rRNA in PWS-IC$^{-p/+m}$ mice. Gene expression was analyzed by RT-qPCR in brains of newborn PWS-IC$^{-p/+m}$ mice. Gapdh was used for normalization. Each data point represents 1 animal (N=6 per genotype), plotted as mean ± SD. Differences to the unaffected controls were analyzed with a 2-tailed, type 3 (assumes unequal variance) Student's t test. *p < 0.05.

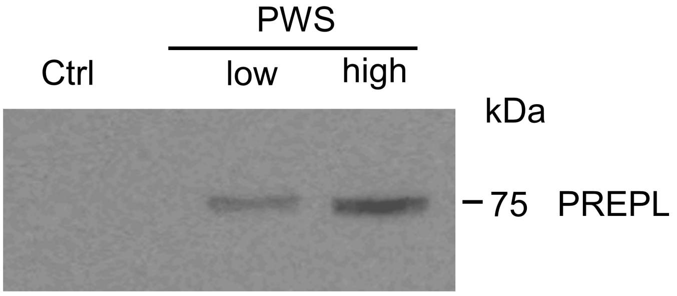


**Supplementary figure 6:** Validation of PREPL expression in fibroblast cell lines. Control (Ctrl) cells transduced with Lenti-GFP shows no expression of PREPL.

### *Genotyping*

| Primer name | Mouse model | Forward | Reverse |
|---|---|---|---|
| **Δ** | *Prepl*$^{-/-}$ | TCTTGCTGTTCCTCCTAGCC | GTCCTGACAAACGGAAAAGG |
| **LoxP** | *Prepl*$^{-/-}$ | GGCAGCTGTAGGAAGTCAGC | ATGTCACAGGCTCGTGTTTG |

| Primer name | Mouse model | Sequence |
|---|---|---|
| **Intron 5** | PWS-IC$^{-p/+m}$ | CCG CAT TTC ATC ATT CTC AGG CTC |
| **PGKneo22** | PWS-IC$^{-p/+m}$ | GTC ACG TCC TGC ACG ACG CGA G |

| Mouse model | Sequence |
|---|---|
| Snord116$^{-p/+m}$ | AAT CCC CAA CCT ACT TCA AAC AGT C |
| Snord116$^{-p/+m}$ | TGG ATC TCT CCT TGC TTG TTT TCT C |
| Snord116$^{-p/+m}$ | TTT ACG GTA CAT GAC AGC ACT CAA G |

**Supplementary table 1:** Primer sequences used for genotyping of the mice (listed in 5'-3' direction). Related to methods.

| ID | Forward | Reverse |
|---|---|---|
| **Guide seq** | CACCGTTACCAAGAAGGTTGTTGCT | AAACAGCAACAACCTTCTTGGTAAC |
| **Sanger seq** | TGACTCCCACTGTACAGGTATCA | CTAGCATAGTACCCAAGACACAGC |

**Supplementary table 2:** Primers for CRISPR-Cas9 mediated HEK293T cloning and sequencing.

| ID | Forward | Reverse |
|---|---|---|
| **Guide seq** | CACCGTAGTTTACCTTCTTCGTCTT | AAACAAGACGAAGAAGGTAAACTAC |
| **Sanger seq** | GAAAGGATACCAGAATGTTG | ACCTGACACACGTAACATATG |

**Supplementary table 3:** Primers for CRISPR-Cas9 mediated βTC3 cloning and sequencing.

### *RT-qPCR*

| Genes | Forward | Reverse |
|---|---|---|
| ***mPrepl*** | TTCTAAGCCAGAGCTCCTGAGAG<br>AACAGAGTTACCGTAACGTC<br>GTCGTTTCCTCACACTGAATATTA | TCCAAGAAAGGTGCCTCCAG<br>TCACTTTCTGGCTTAATAGGCA<br>GCTTGAATTCTGTAGGTTCTCC |
| ***mNhlh2*** | GTGTCGGACCTAGAGCCAGT | CAGGTTGAAAGCCTCCACTC |
| ***mPcsk1*** | TGGAGTTGCATATAATTCCAAAGTT | AGCCTCAATGGCATCAGTTAC |
| ***mPomc*** | CCATAGATGTGTGGAGCTGGT | CAGTCAGGGGCTGTTCATCT |
| ***mNpy*** | TGGACTGACCCTCGCTCTAT | TGTCTCAGGGCTGGATCTCT |
| ***mAgrp*** | TGTGTTCTGCTGTTGGCACT | ACTTCTTCTGCTCGGTCTGC |
| ***mGhsr1a*** | GACCAGAACCACAAACAGACAG | GGCTCGAAAGACTTGGAAAA |
| ***mGapdh*** | CCCCAATGTGTCCGTCGTG | GCCTGCTTCACCACCTTCT |
| ***mSnrpn*** | CTGAGGAGTGATTTGCAACGC | CAAGATCCTTAATACTCGGGG |
| ***mSnord116*** | TGGATCTATGATGATTCCCAG<br>TGGATCTATGATGATTCCCAG | TGGACCTCAGTTCCGATGAG<br>TGGACCTCAGTTCCGATGAG |
| ***mNd1*** | TGCACCTACCCTATCACTC | ATTGTTTGGGCTACGGCTC |
| ***mNdufs3*** | TGTCTCTGCGGTTCAACTCT | GGATGTCCCTCGAAGCCATA |
| ***mCytb*** | TACCTGCCCCATCCAACATT | TAAGCCTCGTCCGACATGAA |
| ***mSdha*** | ATTTGGTGGACAGAGCCTCA | GGCACTCCCCATTTTCCATC |
| ***mUqcrc1*** | TTGCCCAGAAACACTTGAGC | GTCACGTTGTCTGGGTTAGC |
| ***mCo1*** | ACCCAGATGCTTACACCACA | TGTGATATGGTGGAGGGCAG |
| ***mCox4*** | CCATGTCACGATGCTGTCTG | CTCCCAAATCAGAACGAGCG |
| ***mAtp5*** | GTACTCCGCTCTGATCATCG | CTCTTCTTTTCCTCCGCTGC |
| ***mAtp6*** | CCACACACCAAAAGGACGAA | GAAGGAAGTGGGCAAGTGAG |

**Supplementary table 4:** Primer sequences used for RT-qPCR, (listed in 5’-3’ direction). Related to methods.
